# Head-related transfer function predictions reveal dominant sound transmission mechanisms in a dolphin head

**DOI:** 10.1101/2025.05.22.655550

**Authors:** YeonJoon Cheong, Alexander Ruesch, Matt D. Schalles, Jana M. Kainerstorfer, Barbara Shinn-Cunningham, Wu-Jung Lee

## Abstract

Toothed whales possess specialized anatomical structures in the head, including thin, excavated lower mandible embedded in mandibular fat bodies, complex skull morphology fused with the upper jaw, and extensive air spaces surrounding the middle ears and beneath the skull. In this study, we use finite element modeling to investigate how these structures influence the transmission of water-borne sounds to the ears. The models are based on volumetric representations derived from computed tomography (CT) scans of a live bottlenose dolphin (*Tursiops truncatus*). We iteratively modify the anatomical structures included in the model and use the predicted head-related transfer functions (HRTFs) as a proxy for comparison. Our results show that the mandibular fat bodies, which support a lower sound speed than the surrounding tissues, significantly enhance the forward receiving directionality at echolocation frequencies through refraction, in a manner similar to the melon in shaping the dolphins’ highly directional transmission beams. Additionally, we show that, in the frequencies encompassing dolphin communication signals, the air volumes help block the otherwise complex sound transmission through the bones. These findings highlight convergent evolutionary solutions in toothed whale anatomy to create strong directionality in both sound emission and reception governed by the same physical principles.

## I. Introduction

Echolocating toothed whales, such as dolphins and porpoises, have been the focus of research for decades as an animal model for active sonar sensing (Au, 1993, 2015). They have the remarkable ability to detect and track small targets over long distance (e.g., Au and Snyder, 1980; Ladegaard et al., 2019) and discriminate between minute differences between targets using echolocation (e.g., Au and Pawloski, 1992; Branstetter et al., 2020). The performance of these animals often surpasses that of current human-made sonar systems, particularly in scenarios involving targets of complex shape or internal structures in reverberative environments (Au and Martin, 2012). However, how exactly the toothed whales achieve such performance by orchestrating their unusual anatomical structures in the head and the potentially specialized neural processes in the brain remains an area of intensive investigation.

Toothed whales’ heads have unique anatomical structures surrounding the auditory pathway from the water to the middle ears. Unlike terrestrial mammals, the ear canals of toothed whales are wax-filled, and the outer ears have undergone evolutionary degeneration, such that they no longer are the main pathway for sound reception (McCormick et al., 1970, 1980). The lower mandible is excavated and thinned at the rear ends, forming a contrast with the fused upper jaws and the skulls, where the bones were extensively rearranged through the course of evolution (Roston and Roth, 2019). The rear ends of the lower mandible are embedded in the mandibular fat bodies (MFBs) that extend backward to the middle ears, which are enclosed within a structure known as the tympano-periotic complex (TPC). These anatomical specializations have been hypothesized to allow toothed whales to hear sounds via their lower jaw [the “jaw-hearing theory” (Norris, 1968)], an idea supported by both behavioral and neurophysiological results (Brill et al., 1988; McCormick et al., 1970; Mohl et al., 1999). Imaging technologies such as computed tomography (CT) and magnetic resonance imaging (MRI) scans have revealed porous air-filled fibrous tissues surrounding the TPCs in the extracranial peribullar cavities as well as extensive air volumes beneath the upper jaw-skull complex (Houser et al., 2004; Ketten, 1994, 1997). These structures likely impact sound transmission, given the large impedance differences between air and biological soft tissues, bones, and the surrounding water.

Harnessing the dramatic growth of computing power, finite element modeling (FEM) has been widely used in recent years to better understand how sound propagates from water to the middle ear in toothed whales. Building on the jaw-hearing theory, researches have highlighted the potential importance of the lower mandible, the MFBs, and the various air volumes through a series of investigations using volumetric representations derived from CT scans of multiple toothed whale species, including the common dolphin (*Delphinus delphis*) (Aroyan, 2001; Krysl and Cranford, 2016), Cuvier’s beaked whale (*Ziphius cavirostris*) (Cranford et al., 2008), finless porpoise (*Neophocaena asiaeorientalis sunameri*) (Song et al., 2019), short-finned pilot whale (*Globicephala macrorhynchus*) (Song et al., 2021), and Risso’s dolphin (*Grampus griseus*) (Wei et al., 2024). Early on, the MFBs were proposed to function as “waveguides” to channel sound to the middle ears (Ketten, 1994, 1997). Supporting this hypothesis, models of head-related transfer function (HRTFs, filters describing how sound is transformed by the transmission pathway) predicted dramatic differences in the sound reaching the middle ear depending on whether or not they included MFBs (Aroyan, 2001). Recent time-domain propagation models also confirmed this result (Song et al., 2019, 2021; Wei et al., 2024). Separately, the lower mandible has been proposed to play different key roles in sound transmission, either acting as a “fast acoustic lens” that focuses sound to reach the middle ear (Aroyan, 2001), supporting flexural waves (Cranford et al., 2008), or other mechanisms (Song et al., 2019, 2021). However, a recent study suggested it has limited influence (Wei et al., 2024). Another study showed that the air volumes within the head cause profound changes in the predicted receiving directivity (Aroyan, 2001), supporting the hypothesis that these air volumes help isolate the middle ear from sound passing through the head, thereby preserving clearer interaural differences in support of source localization (Houser et al., 2004). However, the use of multiple toothed whale species across these modeling studies and the sometimes ambiguous interpretation of the underlying physical mechanisms make it challenging to determine if the proposed mechanisms generalize across species or CT scan sources.

Here, we focus on deriving physics-based, mechanistic interpretations of FEM predictions Using a three-dimensional (3D) volumetric representation of the head of a live bottlenose dolphin reconstructed from CT scans. To discover how different anatomical features affect sound transmission, we analyzed how model HRTF predictions change as we iteratively removed or added individual structures, including the MFBs, lower mandible, upper jaw-skull complex, TPCs, and the peribullar and pterygoid air volumes. HRTF analyses are augmented by analytical and numerical models with simple geometries that capture key dimensions and material properties of the measured anatomical structures. This approach differs from most previous studies in which the influence of individual anatomical structures on the HRTF was not explicitly modeled and interpreted based on physics principles. This includes an artificial head volume, assembled using only shapes of simple geometries, which successfully predicts the main HRTF features present in the full model from the head volume reconstructed from CT scans. This modeling approach allows us to interpret the physical underpinnings of dominant sound transmission mechanisms in the toothed whale head, providing important insights that can guide future studies of acoustic transmission in different species, or even different individual animals.

## II. Modeling approaches

### A. Shape representations

A 3D volumetric representation of the head of a bottlenose dolphin was derived from CT scan images of a live bottlenose dolphin that included the tip of the rostrum to the pectoral fins. We segmented the images up to the back of the skull to reconstruct important anatomical structures within the head, including the melon, upper jaw-skull complex, lower mandible, MFBs, TPCs, non-fat soft tissues, and air volumes (Fig. 1A). The MFBs encompass the fat components on both the medial and lateral sides of the jawbone, which were sometimes described as separate structures in previous studies (e.g., Song et al. 2021; Wei et al. 2024). The air volumes included the air in the nasal passages, the fibrous tissues surrounding the TPCs that have a collective density very similar to air (hereafter the “peribullar air volumes”), and the pterygoid air volumes located beneath the upper jaw-skull complex (Houser et al., 2004). The segmentation was performed manually using 3D Slicer (Fedorov et al., 2012) with the following Hounsfield Unit (HU) ranges: bone: (200, 2143), fat: (-150, 0), air: (-1000, -500). The non-fat soft tissues were obtained by segmenting the CT scan images using HU values larger than -300 and excluding all other structures. These ranges were selected based on previously reported values (Song et al., 2017). Additional smoothing of each segmented component was performed to allow successful meshing of the computing domain in COMSOL Multiphysics.

**Figure 1.**
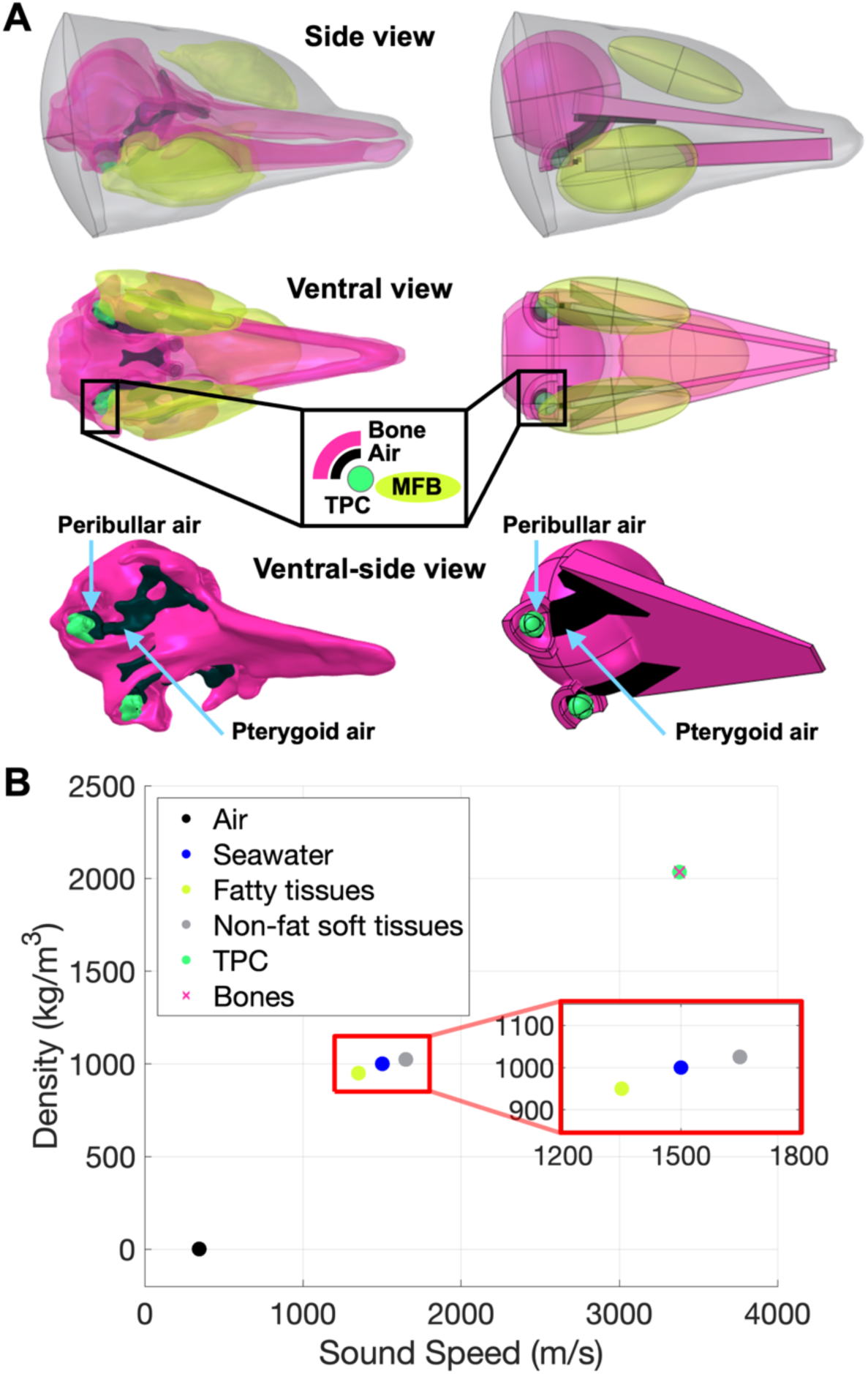
(**A**) 3D volumetric representation of the head of a bottlenose dolphin with key anatomical structures color-coded according to their material property categories. Left: CT-derived full-head model; Right: Artificial-head model. (**B**) Sound speed and density for different material categories, including the surrounding seawater as reference. Note the TPCs are assigned the same material properties as other bones, but they are plotted separately for clarity.

The material properties of the segmented structures (Fig. 1B) were assigned by referencing experimentally measured values reported in previous studies. For soft tissues, we estimated the midpoints of the values reported in Song et al. (2017): Fatty tissues, including the melon and the MFBs, were modeled as fluid, with a pressure wave speed of 1350 *m*/*s* and a density of 950 *kg*/*m*^3^. Non-fat soft tissues (flesh) were also modeled as fluid, with a pressure wave speed of 1650 *m*/*s* and a density of 1025 *kg*/*m*^3^. Bones, including the upper jaw, the lower mandible, and the TPCs, were modeled as elastic solids with a pressure wave speed of 3380 *m*/*s*^3^, a shear wave speed of 2200 *m*/*s*, and a density of 2035 *kg*/*m*^3^ using values from Dible et al. (2009) that were used in Song et al. (2019, 2021).

To help interpret our results we also constructed a 3D volumetric head representation using shapes of simple geometries that mimicked the primary dimensions and shapes of the actual structures (hereafter “the artificial-head model,” Fig. 1A right column). Specifically, the MFBs, lower mandible, upper jaw, peribullar air volumes, and skull were modeled as ellipsoids, rectangular beams, pyramids, quadrants of spherical shells with a thickness of 1 mm (see inset of Fig. 1A), and a spherical shell with the inner sphere offset toward the posterior direction, respectively. The pterygoid air volumes are modeled by combining a 5 mm-thick pyramid-shaped plate that represents the air volume beneath the upper jaw and a 3 mm-thick partial spherical shell that represents the air volume beneath the skull (see the ventral view in Fig. 1A). We used this artificial-head model to understand how the major features in the HRTF patterns computed from the full, realistic head model relate to the anatomical features in the specific CT scans and to make sure that the full-model results are not idiosyncratic or due to peculiarities arising from the segmentation procedure.

### B. Numerical modeling software and computing resources

All numerical simulations were performed using COMSOL Multiphysics (version 6.1) solvers for coupled acoustics and solid mechanics equations to account for the combined effects of the various anatomical structures in the dolphin head. The acoustic-structure interaction interface was used to solve the coupled equations and ensure computational stability. The computational domains, including the surrounding seawater medium and the structures in the head, were meshed with mesh sizes no larger than one fifth of the wavelength at each of the frequencies analyzed. We have benchmarked the accuracy of the models computed using this criterion by comparing the predicted target strength of elastic spheres and spherical shells of different material properties and sizes with the exact modal series solution (Faran, 1951).

Given the large domain enclosing the dolphin head (a sphere with a diameter of 54 cm, Fig. 2), computing full-wave simulations at high frequencies with corresponding short wavelengths (max. 50 kHz with a wavelength of approx. 0.03 m in this work) requires substantial memory resources due to the fine mesh size requirements.

**Figure 2.**
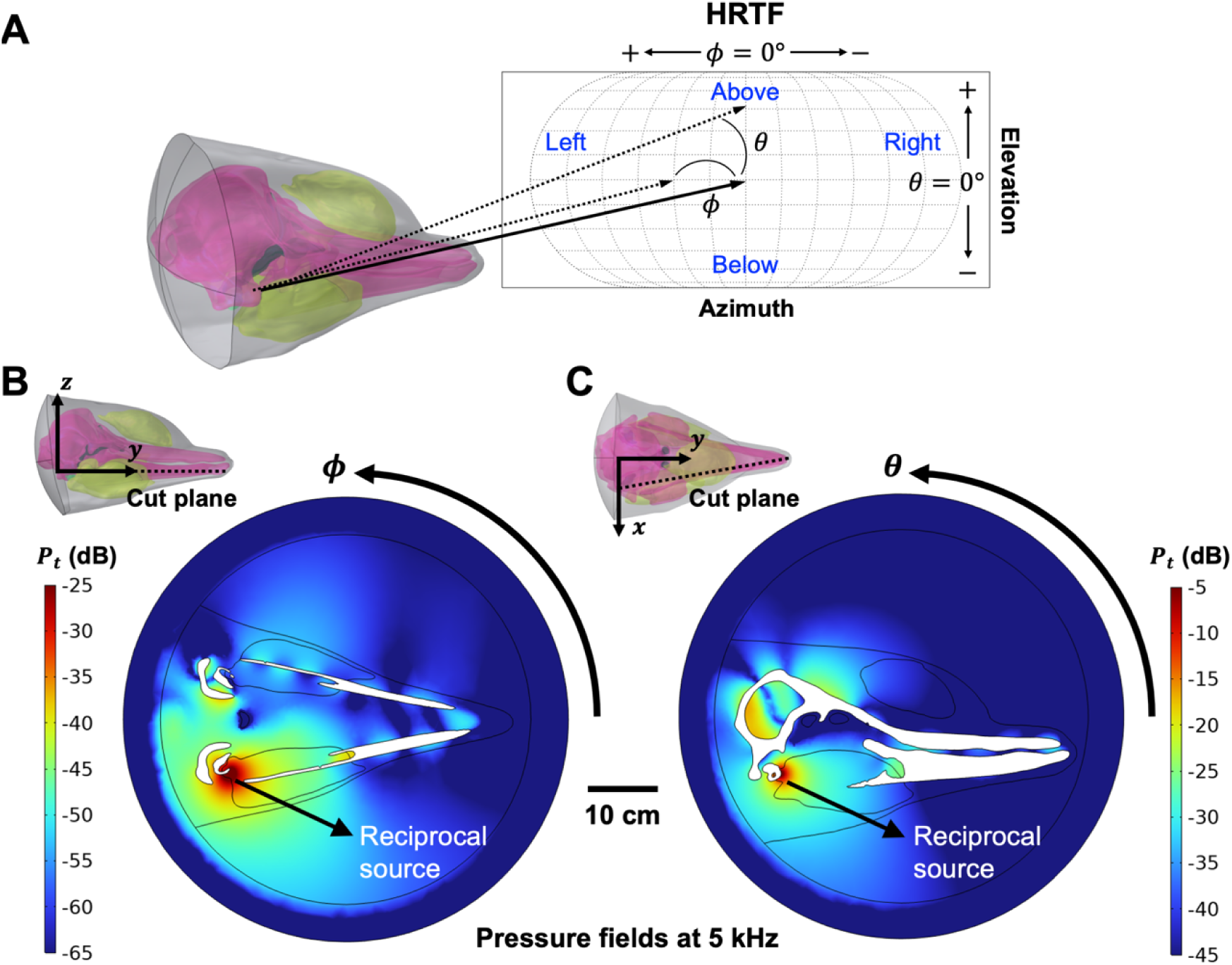
(**A**) A schematic showing the dolphin-centric polar coordinate scheme, where the HRTF is a function of azimuth (ϕ) and elevation (θ). The origin of the coordinate is at the midpoint between the left and right “ears” near the TPCs (see Sec. II.C for detail). (**B-C**) Example total pressure fields computed using the full-head model at 5 kHz on a horizontal and a vertical plane. The horizontal plane cuts through the left and right ears and the lower mandible (i.e., this plane includes the origin). The vertical plane is perpendicular to the horizontal plane and cuts through the right ear and the tip of the rostrum.

### C. Head-related transfer functions (HRTFs)

The HRTF is a mathematical description of how sound originating from an external source at azimuth ϕϕ and elevation θθ relative to the middle ear is transformed by different anatomical structures (Fig. 2A). The HRTF is defined as (Gumerov et al., 2010)

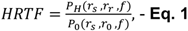

where *r_s_* denotes the source position, *r*_r_ denotes the ear position at the end of the MFBs at the vicinity of the TPC (with *r_r,L_*_,*_ and *r_r,R_* denoting the left and right ears, respectively), *P_H_* is the total sound pressure at *r_r_* with the head present, *P*_0_ is the total sound pressure in the water domain without the head at the midpoint between the two ears (i.e., *r*_0_ = (*r_R, L_*, + *r_r,R_*)/2), and *f* is the sound frequency. HRTF quantifies the ratio of received sound pressure at each ear with and without the presence of the head, and HRTF values>1 and <1 indicate enhancement and reduction of the received pressure at the two ears, respectively. Note that for sounds sufficiently far away from the animal, the sound wave is approximately planar when it impinges on the head and body; in such instances, the effect of distance influences only the overall sound pressure level and does so identically for all frequencies. Thus, for this definition of the HRTF, normalized by the pressure that would be present in free water, the effect of distance in the numerator and denominator cancel, and the HRTF depends only on source direction relative to the animal. Roughly speaking, this assumption of a planar wavefront is reasonable for sounds at distances of 1 m or greater.

We computed the HRTFs for θ from -90° to 90° and ϕ from -180° to 180° (Fig. 2A) at select frequencies between 5 kHz and 50 kHz, spanning the typical range of frequencies important for communication and echolocation. For efficiency, the computation was performed using the reciprocity of sound propagation, with the two ear positions in the non-fat soft tissues near the TPCs (*r_r,L_*_,*_ and *r_r,L_*) being the reciprocal source and the varying sound sources being the reciprocal receivers. The reciprocal receivers were located on a spherical surface at a 2 m distance from the midpoint of the two ears.

Fig. 2B-C show example simulated pressure fields within the spherical computational domain with the reciprocal source located at *r_r, R_* using polar coordinates relative to the horizontal θ=0 plane (Fig. 2B) that cuts through both *r_r, L_* and *r_r, R_* (and therefore *r*_0_) and the lower mandible, and the vertical ϕ=0 plane (Fig. 2C) that cuts through *r_r, R_* and the tip of the rostrum. The sound was propagated beyond the boundary of the computational domain to a distance of 2 m, where the pressure was taken to compute the HRTF.

### D. Investigating the effects of individual anatomical structures

The HRTFs computed using the dolphin head volume contain the combined effects of all segmented anatomical structures (hereafter referred to as the “full-head model”). To investigate the overall influence on the HRTF and the associated sound transmission mechanisms of individual anatomical structures, we first constructed a “base model” consisting of only two TPCs enclosed in non-fat soft tissues with the realistic dolphin head shape, and iteratively constructed a series of “reduced-head models” that each includes one additional anatomical structure on top of the base model (Fig. 3). While the acoustic effects of these individual structures do not combine linearly, this procedure provides a systematic framework to analyze and disentangle complex physical processes. We used the same comparative approach to investigate the effects of incorporating shear waves by modeling the bones as either fluid or elastic solids. We further used shapes of simple geometries to investigate the physical mechanisms associated with each anatomical structure in the head. These simplified geometries and mechanisms are detailed in the following section. Note that the vibroacoustic coupling related to and within the TPCs are outside the scope of this work; we focus exclusively on the transmission of sound from seawater to right before it enters the middle ear. Additionally, reduced-head models with the nasal air volume or the melon added were computed but not included here, because they did not contribute substantially to the HRTF.

**Figure 3.**
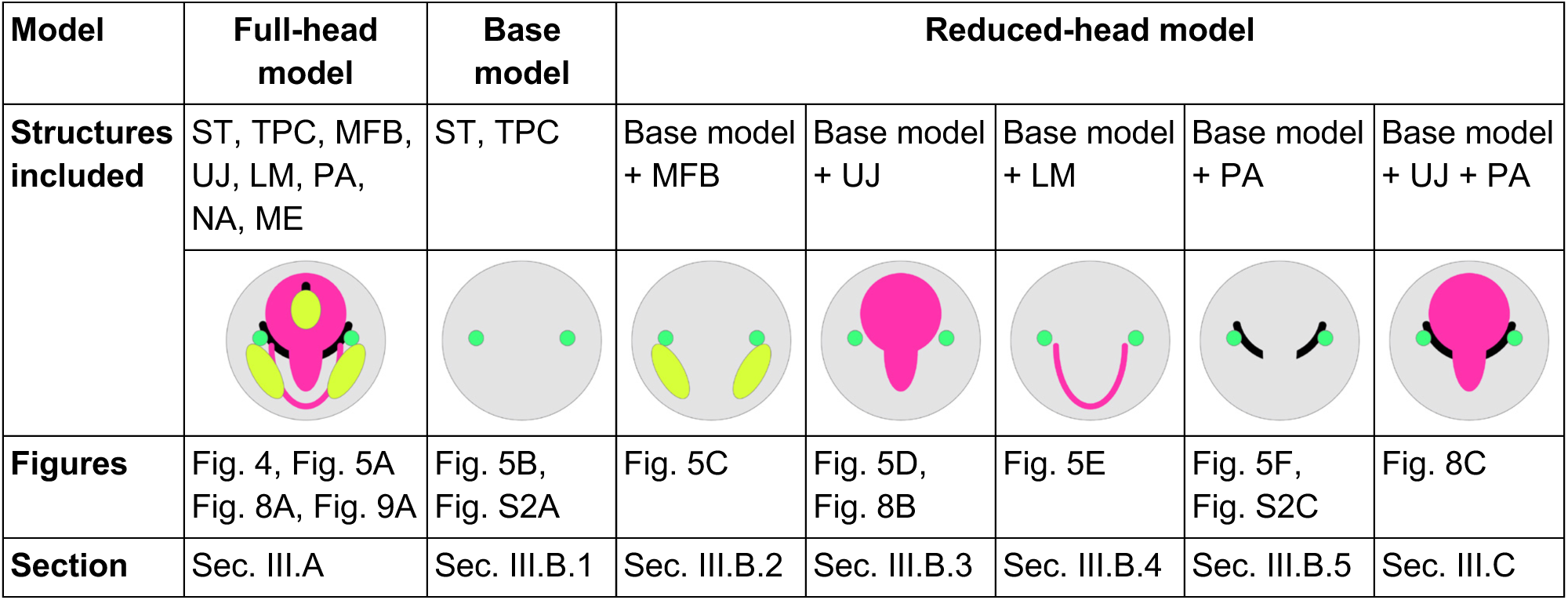
Schematics illustrating the different anatomical structures included in the volumetric representations used to compute HRTFs for the full-head model and the various comparison models. Note that to make the schematics easy to interpret, the sizes of the structures are not drawn to scale and their relative positions are not accurately represented. LM: lower mandible; ME: melon; MFB: mandibular fat body; NA: nasal air volume; PA: peribullar and pterygoid air volumes; ST: non-fat soft tissues; TPC: tympano-periotic complex; UJ: upper jaw-skull complex.

## III. Modeling results

### A. HRTFs of the full-head model

The HRTFs computed using the full-head model show strong direction-dependent features that vary across frequency (Fig. 4). Specifically, the full-head HRTFs show broad pressure enhancement regions (sensitive regions) for sounds arriving from a direction ipsilateral to the receiver (e.g., the left receiver is more sensitive to sound originating from the left-hand side) at lower frequencies. These sensitive regions shrink in size and shift toward the front of the animal with increasing frequency. Note the HRTFs outside the azimuth angular range of ϕ=-90°-90° may not be reliable as our model did not include any structures in the dolphin body behind the skull.

**Figure 4.**
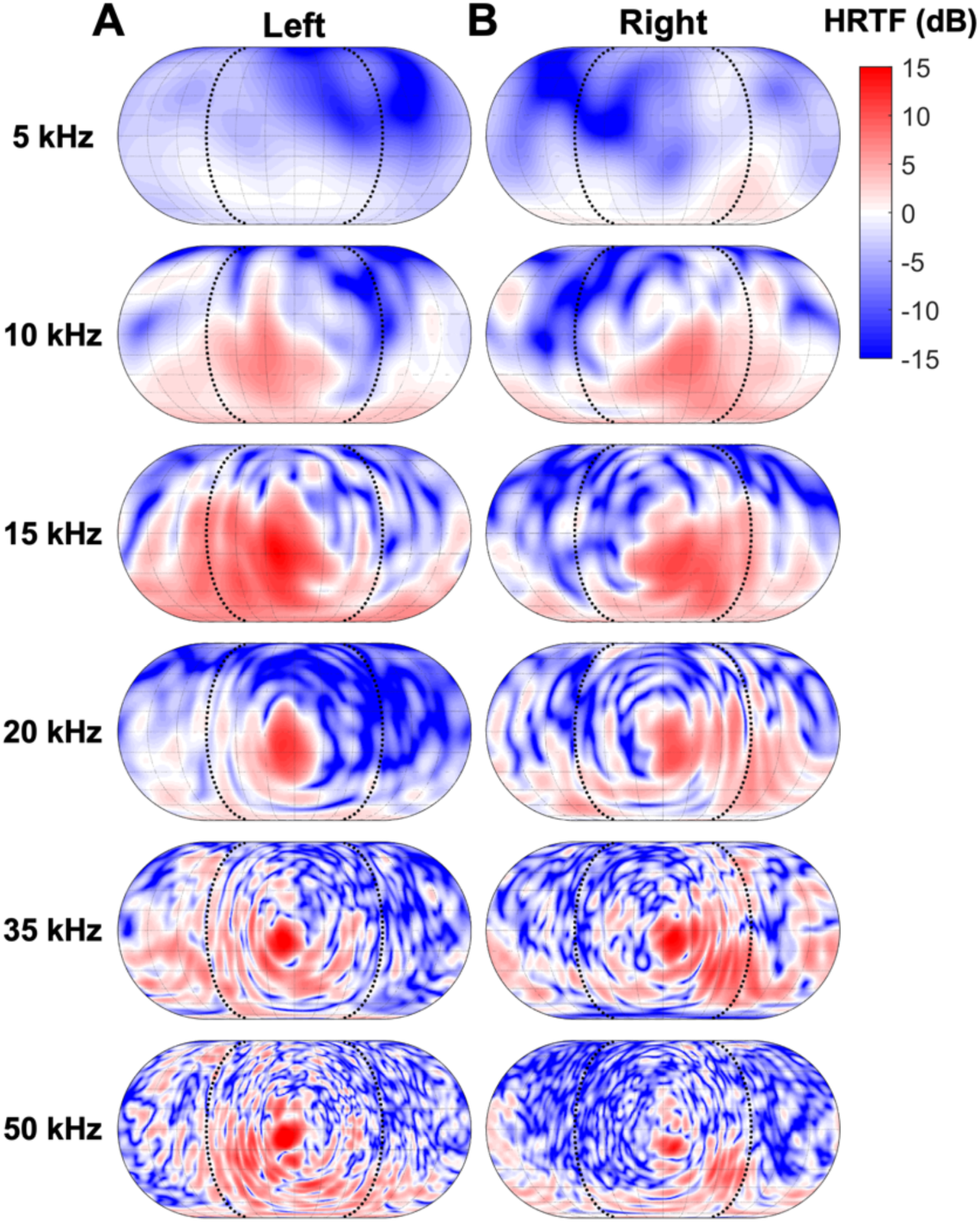
HRTFs at the left and right ears computed using the full-head model. Within each panel, the gain (red) or reduction (blue) in the received sound level compared to open water is plotted as a function of the direction of the sound source relative to the ear. Each row shows results for a different frequency, ranging from 5 kHz (top) to 50 kHz (bottom), with results for the left ear in the left column and the right ear in the right column. The black dotted lines indicate the reliable azimuthal range between ϕϕ = -90° to 90°, since the model did not include any structures in the dolphin body behind the skull. The HRTFs are left-right symmetric in their broad features but show asymmetry in their detailed structure. The directions of sound producing the greatest received energy form large sensitive regions (red areas in the plots) at low frequencies that shrink in size and shift toward the front of the animal with increasing frequency.

The predicted HRTF patterns are broadly left-right symmetric, with a moderate asymmetry that likely is due to the broader left nasal passage on the skull (Coombs et al., 2020; Cranford et al., 1996). Similar asymmetric HRTF patterns have been reported previously from common dolphins (Aroyan, 2001; Krysl and Cranford, 2016). Details in the HRTF patterns also are influenced by anatomical variability and idiosyncrasies across individuals, especially the exact shapes and sizes of air cavities in the head measured during the CT scan, due to the large impedance differences between air and body tissues and bones. While the left-right asymmetry may be important in supporting source localization as shown in other animals (e.g., Knudsen and Konishi, 1979), systematic comparisons of HRTFs predicted using CT images across multiple individuals of the same and different species within a phylogenetic framework are likely needed to address this question, which is beyond the scope of this present study.

### B. Influence of different anatomical structures

In this section, we compare the HRTFs computed using the full-head model, the base model, and a series of reduced-head models to illuminate how different anatomical structures influence the overall HRTF patterns. Recall that the base model consisted of only the non-fat soft tissues and the TPCs, and the successive reduced-head models each include one additional anatomical structure on top of the base model (Fig. 3). Importantly, we reiterate that the various anatomical structures interact nonlinearly, not through superposition, in the full-head model; the comparisons with reduced-head models simply provides a way to gain insight into the complex sound transmission processes involving multiple structures.

#### 1. Base model

The HRTFs computed using the base model (Fig. 5B) show a prominent frontal region of pressure reduction, with the size shrinking at higher frequencies. This can be explained intuitively by the higher sound speed of seawater compared with the non-fat soft tissues (Fig. 1B), which causes the sound to refract away from the center of the head. This is the reverse of the effects of the MFBs (see Sec. III.B.2 below). The HRTFs from the base model provide another reference point along with the full-head model for comparisons with the series of reduced-head models discussed below.

**Figure 5.**
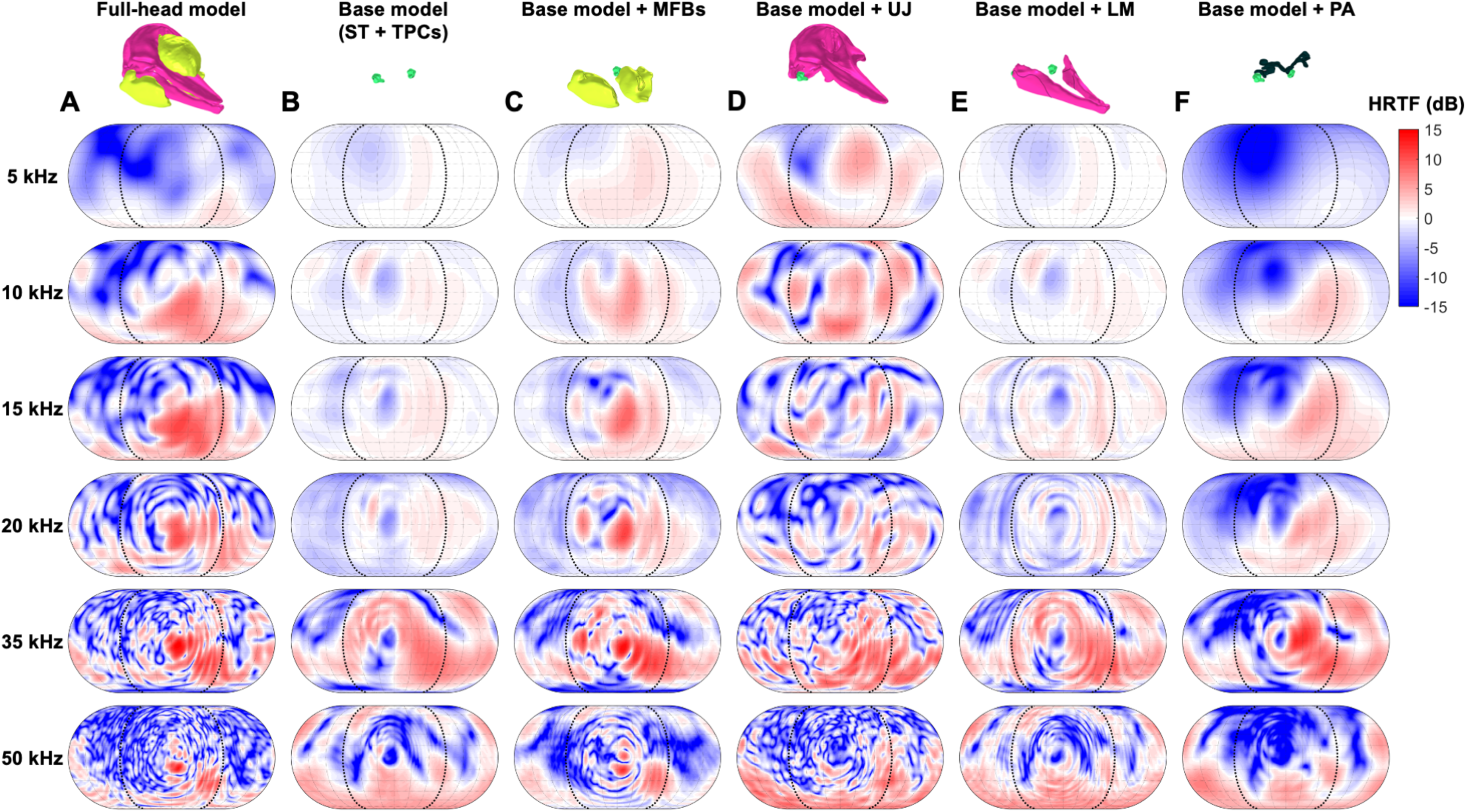
HRTFs at the right ear computed using the (**A**) full-head model and (**B-F**) a series of reduced-head models. The HRTFs are plotted using the same color scale as in Fig. 4. See Fig. S3-S4 for corresponding HRTFs computed at the left ear.

The HRTFs also exhibit larger swaths of pressure enhancement at higher frequencies (>35 kHz). Based on further investigation of the scattering characteristics of the TPC (not shown), these features are likely due to vibroacoustic coupling occurring on a single TPC and the interaction between the two TPCs, which are beyond the scope of this work.

#### 2. Mandibular fat bodies

The HRTFs computed using a reduced-head model consisting of the non-fat soft tissues, the TPCs, and the MFBs (Fig. 5C) show a prominent pressure enhancement region in front of the animal at azimuth angle between -30° and 30°, with the size of the region shrinking with increasing frequency. Around this region, the HRTFs broadly resemble the patterns from the full-head model (Fig. 5A) at ≥15 kHz, and across most frequencies show gain patterns that are almost opposite of the patterns obtained from the base model (Fig. 5B). The HRTF patterns can be explained by the focusing effects caused by the refraction of sound through the MFBs, which have a smaller sound speed than the non-fat soft tissues and the seawater (Fig. 1B).

To better understand this interaction, we computed the equivalent of HRTFs from a reciprocal source located right next to the apex of an ellipsoid with the same material properties and dimensions similar to the MFBs (Fig. 6). The slower sound speed within the ellipsoid and its convex shape jointly form a classic acoustic lens, through which waves with collimated wavefronts are refracted and focused to a single point at the high frequency limit (Beaver et al., 1977). Here, the narrowing main lobe and the frequency-dependent changes of the sidelobes (decreasing spacing with increasing frequency) are similar to the patterns from the reduced-head model (Fig. 5C).

**Figure 6.**
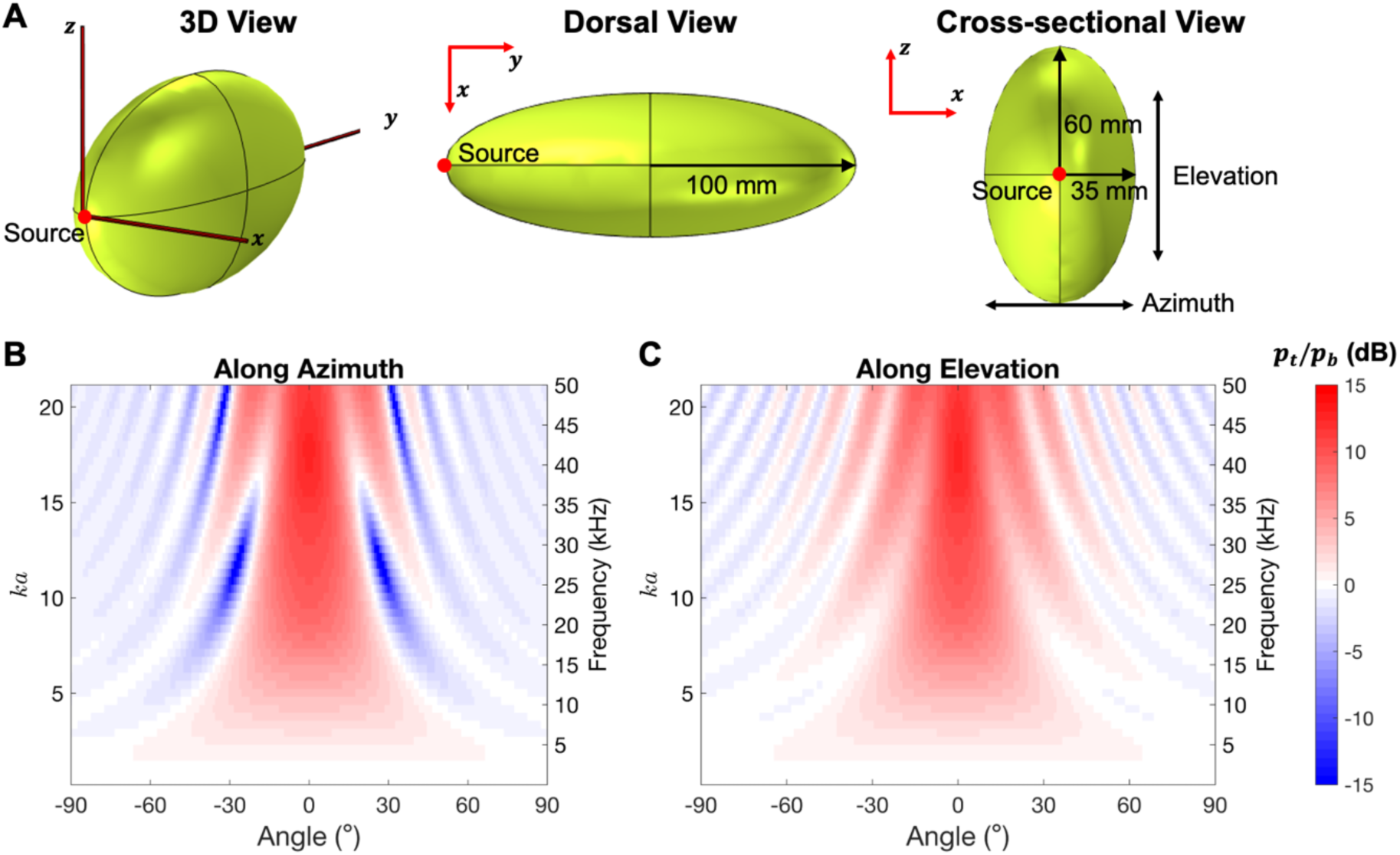
(**A**) A mandibular fat body (MFB) modeled as an ellipsoid shown in three perspectives. (**B-C**) The equivalent of HRTF computed using a reciprocal receiver located right next to the apex of the ellipsoid. The largest half length of the ellipsoid semi-axes (*a*=100 mm) is used to calculate the *kga* values in panels B-C.

#### 3. The upper jaw-skull complex

The HRTFs computed using a reduced-head model comprising only the non-fat soft tissues, the TPCs, and the upper jaw-skull complex (Fig. 5D) show prominent sensitive regions at 5 kHz and 10 kHz and more fragmented patterns at higher frequencies. The general HRTF patterns are quite different from those from the full-head model (Fig. 5A), except for the pressure enhancement regions at 10 kHz. Specifically, the HRTFs lack the frontal pressure enhancement regions at higher frequencies due to the MFBs (see Sec. III.B.2). Compared to the base model, the HRTFs show dramatic enhancement at broad angular ranges, particularly at lower frequencies (up to 20 kHz), and more fragmented patterns with the sensitive regions displaced toward the right at 35 kHz and 50 kHz.

To better understand the mechanisms underlying these HTRF patterns, we constructed a simple 2D model composed of a bone layer sandwiched in two half-spaces filled with non-fat soft tissues (Fig. 7). The bone layer was modeled either as a fluid (Fig. 7A), which supports only pressure waves, or as an elastic solid (Fig. 7B), which supports both pressure and shear waves. For sound transmitted through the bones, the transmission coefficient is *T*1_13_ =↓ *P*1_3_/↓ *P*_1_, where ↓ *P*1_1_ is the incident pressure wave in the upper half-space and ↓ *P*_3_ is the transmitted pressure waves in the lower half-space. *T*1_13_ is computed for all incident angles (*α*_1_) and a range of *k*_1_*h,* where *kg*_1_ is the wavenumber in medium 1 and ℎ is the thickness of the bone layer. Detailed derivation and analytical expressions for these quantities are provided in the Supplemental Text S1.

**Figure 7.**
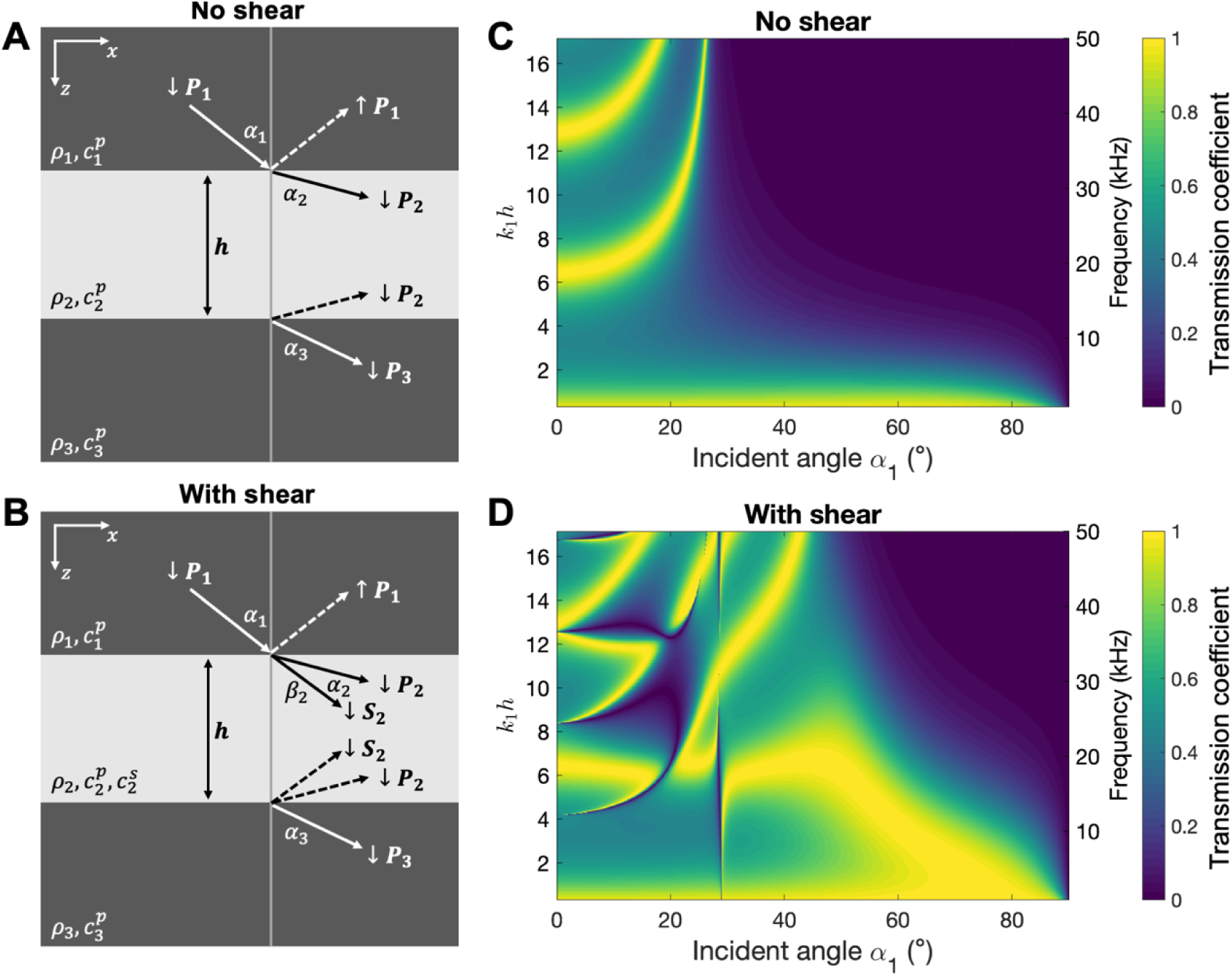
(**A-B**) Schematics of 2D models with a bone layer sandwiched between non-fat soft tissues. (**C-D**) The transmission coefficients across the bone layer. The bone layer is modeled as either fluid (panels A and C) or an elastic solid (panels B and D). The *k*_1_ℎ values are computed using a fixed bone layer thickness ℎ=90 mm, which is roughly the dimension of the thickest section of the upper jaw.

When the bone layer is modeled as fluid (Fig. 7A and 7C),*T*_13_ is zero or very low for the majority of angles, except for the scenarios where *k*_1_ ℎ ≈ 0 and along the branches at ℎ = 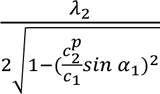, where λ_2_ is the pressure wavelength in the bone layer. Here, α*_c_* = 29.22^∘^ is the critical angle beyond which total reflection occurs. When the bone layer is modeled as an elastic solid (Fig. 7B and 7D), there are additional broad high-transmission regions above α*_c_* at *k*_1_ℎ < 7 and additional branches at higher*k*_1_ ℎ, as well as additional low-transmission regions below α_c_ for *k*_1_ℎ = 5 − 15. The presence of these high-transmission regions and branches is consistent with the broader sensitive regions at frequencies <10 kHz and the more fragmented pattern of sensitive regions at >15 kHz shown in the HRTFs from the reduced-head model (Fig. 5D). The overall effects of incorporating shear waves on the HRTFs will be discussed further in Sec. III.C.

#### 4. Lower mandible

The HRTFs computed using a reduced-head model consisting of only the non-fat soft tissues, the TPCs, and the lower mandible (Fig. 5E) bear little resemblance to the HRTFs computed using the full-head model (Fig. 5A). Compared with the base model (Fig. 5B), the addition of the lower mandible appears to introduce interference patterns to the total sound field, which alter the detailed striation or induce additional striation patterns in the HRTFs.

At lower frequencies, the small contribution of the lower mandible to the overall HRTF patterns is at least in part related to the small transverse dimension of these thin bones, particularly in the posterior “pan bone” area. This falls in the small *kgkg*ℎ regions in Fig. 7D, at which sound largely transmits effectively through the bones. At higher frequencies, it is possible that the thin and broad shape of the lower mandible lead to the formation of flexural waves (the zeroth-order asymmetrical lamb waves) (Cranford et al., 2008) or other surface waves that propagate at a different speed than the pressure and shear waves, thereby causing complex interference patterns in the HTRFs.

#### 5. Peribullar and pterygoid air volumes

The HRTFs computed using the reduced-head model that includes the non-fat soft tissues, the TPCs, and the peribullar and pterygoid air volumes (Fig. 5F) show prominent pressure reduction for sound originating from the contralateral and dorsal-frontal side of the receiver (e.g., sounds from the left-hand side and the forward directions are not conducted effectively to the right ear). The dramatic difference between these HRTFs and those computed using the base model (Fig. 5B) demonstrates that the pressure reduction is the result of the presence of air volumes, as this is the only difference between these two models. The HRTF patterns are also dramatically different from those computed using the full-head model (Fig. 5A), but the combined influence of other structures are more difficult to isolate.

The pressure reduction arising from the air volumes is likely explained by strong reflections off the interface between air and the non-fat soft tissues or the bones, which effectively blocks sound transmission. Indeed, the transmission coefficient *T*≈ 0 for all incident angles in a simple 2D model consisting of a half-space with material properties of the non-fat soft tissues or the bones and a half-space filled with air [Supplemental Text S1, eq. (6)].

### C. Effects of incorporating shear waves

To better understand the importance of incorporating the effects of shear waves, we compared HRTFs while modeling the bones as either an elastic solid or a fluid using 1) the full-head model, which includes all structures, 2) a reduced-head model consisting of only the TPCs, non-fat soft tissues, and the upper jaw-skull complex, and 3) a reduced-head model consisting of all structures in 2) as well as the peribullar and pterygoid air volumes.

For HRTFs computed using the full-head model (Fig. 8A), the effects of incorporating shear waves manifest primarily as additional pressure enhancement regions. These regions appear as larger branches with moderate enhancements at lower frequencies (<15 kHz) and numerous fragmented small regions at higher frequencies (≥15kHz). In contrast, the HRTFs computed without shear wave effects appear “cleaner,” with larger, continuous pressure reduction regions in the contralateral direction.

**Figure 8.**
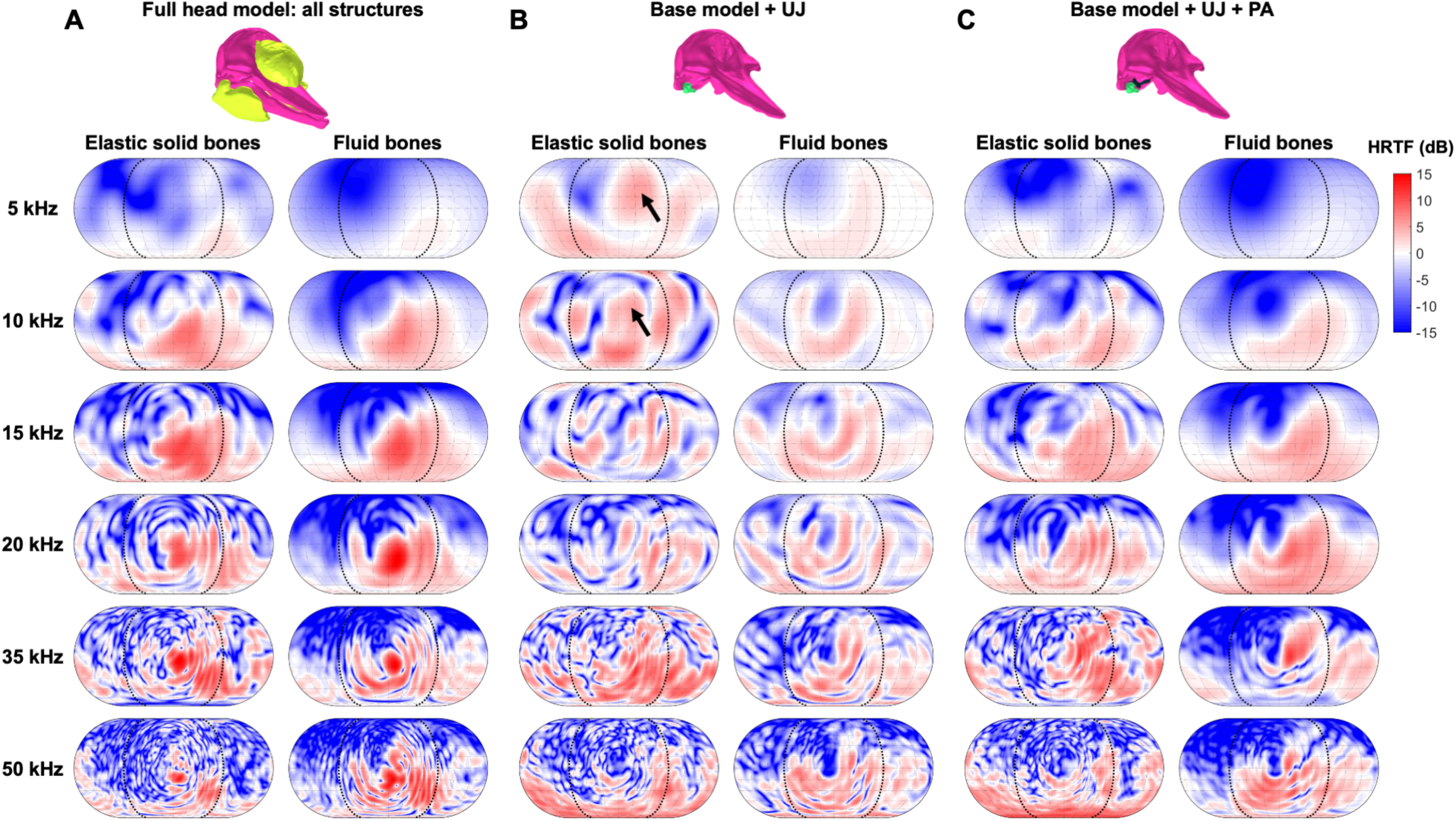
Comparison of HRTFs at the right ear computed by modeling bones as either fluid or elastic solids. (**A**) The full-head model. (**B**) A reduced-head model consisting of the TPCs, non-fat soft tissues, and the upper jaw-skull complex. (**C**) A reduced-head model consisting of all structures in panel B as well as the peribullar and pterygoid air volumes. Note the left columns in panels A, B showing elastic solid results are identical to Fig. 5A and Fig. 5D, respectively. See Fig. S3-S5 for corresponding HRTFs computed at the left ear.

The HRTFs computed using the reduced-head model without the air volumes (Fig. 8B) show that the difference observed for the full-head model can be attributed to the additional sound transmission paths through the bony upper jaw-skull complex from the dorsal-frontal direction (indicated by arrows in Fig. 8B) when the bones are modeled as an elastic solid. This is consistent with the larger swath and additional branches with high transmission coefficients shown in Fig. 7D when the bone layer in the 2D model was modeled as an elastic solid.

Interestingly, including the peribullar and pterygoid air volumes in the reduced-head model reduces these differences (Fig. 8C), likely because they collectively form an effective barrier of sound originating from the dorsal-frontal and contralateral directions (see Sec. III.B.5).

### D. HRTFs of an artificial-head model built from simple shapes

To qualitatively validate the above simulations and interpretations against potential idiosyncrasies associated with the particular CT scan images of a single dolphin, we constructed an artificial-head model from a handful of shapes with simple geometries, each capturing an anatomical structure segmented from the CT images (see Sec. II.A for detail).

The overall HRTF patterns computed using the full-head model and the artificial-head model are broadly similar (Fig. 9). Refraction through the MFBs, which gives rise to the strong frontal sensitive region, appears to consistently dominate the HRTFs at the echolocation frequency range (≥ 15 kHz). The broad pressure enhancement and reduction regions at lower frequencies and the complex striations at higher frequencies are also reproduced in the HRTFs computed using the artificial-head model, although the exact locations and angular extents of these regions differ. These differences were expected, since the combined results of sound transmission through the upper jaw-skull complex and reflection from the air volumes are sensitive to the specific geometry and angles between these structures and the sound source direction.

**Figure 9.**
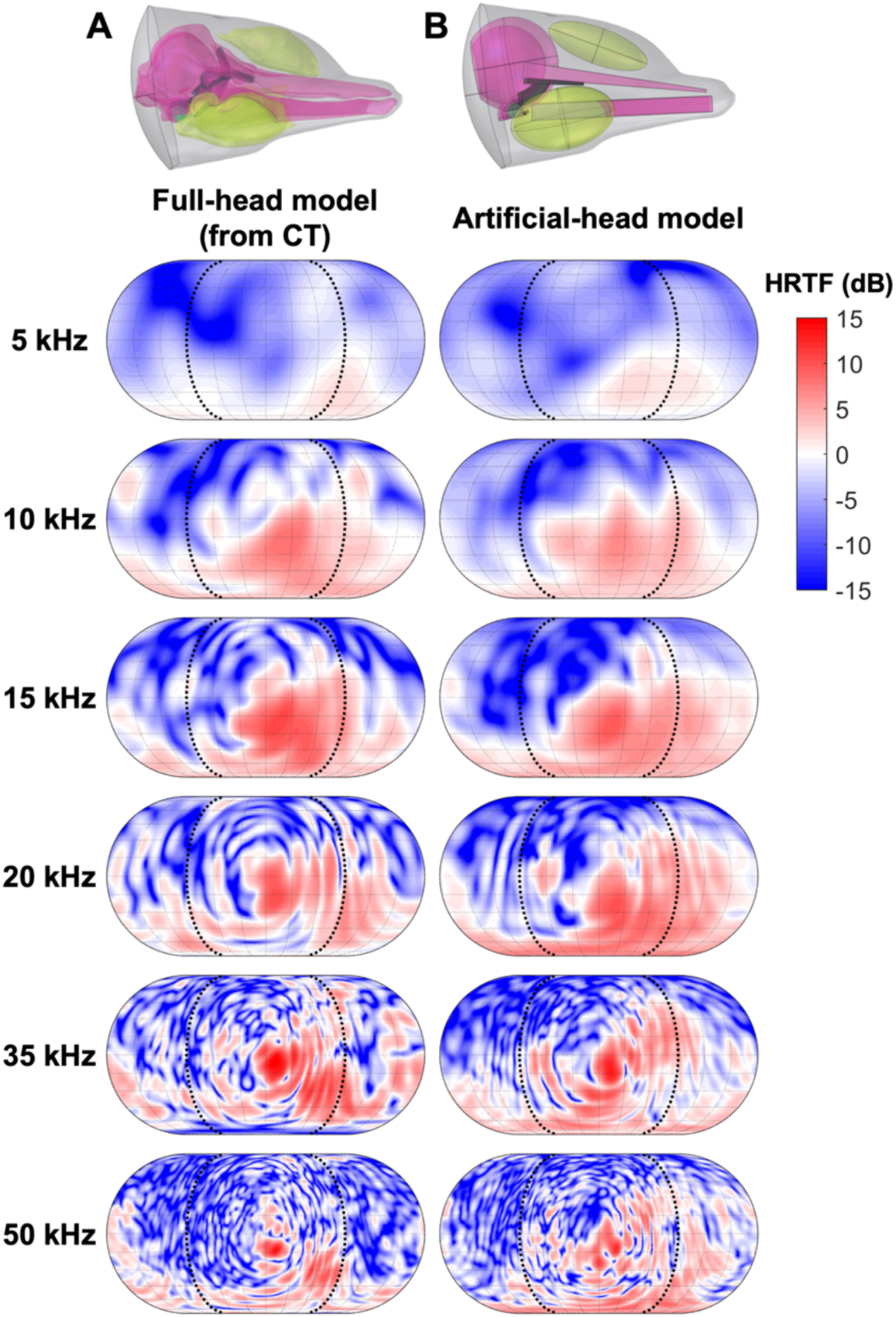
HRTFs at the right ear computed using the full-head model derived from CT scans (left column) and the artificial-head model constructed using shapes of simple geometries (right column). See Sec. II. A for details. See Fig. 4 and Fig. S6 for HRTFs at the left ear computed using the full-head model and the artificial-head model, respectively.

Note that the HRTFs computed by the artificial-head model are left-right symmetric due to the absence of asymmetry in the artificial head volume. We did not introduce asymmetries in the artificial head volume, because our goal was to illustrate that matching gross dimensions and shapes is sufficient to reproduce the dominant HRTF patterns from CT-derived head volume. This approach highlights the underlying physical mechanisms of sound transmission, rather than to precisely replicate every detail of the HRTFs.

## IV. Discussion

In this study, we investigated how sound propagates from the water to the middle ear in the toothed whale head by analyzing model HRTFs computed using FEM. Our goal was to provide a physics-based mechanistic understanding of the sound transmission processes. To elucidate the physical underpinning of the observed HRTF patterns, the models used both realistic 3D volumetric representations of the anatomical structures in a dolphin head as well as artificially constructed structures with simple geometrical shapes. Our modeling results show the critical role of the MFBs in focusing sound from the forward direction in the echolocation frequency range. They also demonstrate the key role air volumes within the head play to block and isolate sound transmission through the skull to the ears, especially at lower frequencies. These findings corroborate and extend previous experimental and modeling results, which we discuss below.

Our model results broadly support the jaw-hearing theory that sound is transmitted to the TPCs via the MFBs (Norris, 1968), but specifically show that the MFBs act as acoustic lenses to guide sound arriving from the front of the animal toward the TPCs (Fig. 5C and Fig. 6), especially for signals falling in the echolocation frequency band (≳15 kHz). This lensing mechanism operates analogously to the way that the melon, another fatty body located in the forehead of toothed whales, is thought to function in conjunction with the skull and the vestibular air sacs to focus outgoing echolocation clicks to form a directional beam (Aroyan et al., 1992; Cranford et al., 2014; McKenna et al., 2012; Song et al., 2021; Wei et al., 2016, 2017). Prior work has described the MFBs as waveguides that enhance sound propagation from the water to the dolphin ears (Ketten, 1994, 1997). Indeed, there is a dramatic enhancement of acoustic energy density “passing through and below the pan bones and extending back to the region of the ear complexes” at 50 kHz (Aroyan, 2001). In the current work, adding the MFBs and other soft tissues produces a directional sensitivity favoring sources from in front of an animal. These results are consistent with the observation that removing MFBs from the model volume decreases sound pressure level at the surface of periotic bones (Wei et al., 2024), and do not contradict with the proposed “gular” pathway where sound also enters the head via the throat region below the lower jaw (Cranford et al., 2008; Wei et al., 2024). Our models help isolate the effects of non-fat soft tissues and the MFBs (Fig. 5B vs Fig. 5C) and show the dominant waveguiding role of the MFBs through a focusing mechanism (Fig. 6).

Our modeling results are also consistent with the hypothesis that the peribullar and pterygoid air volumes function as sound-reflecting barriers (Aroyan, 2001) and contribute to the animals’ ability to differentiate the direction of incoming sound and damping reception of self-generated sounds (Houser et al., 2004). The HRTFs computed with and without these air volumes (Fig. 5F vs Fig. 5B, respectively) show their effects in blocking sound from the dorsal-frontal and contralateral directions. The peribullar air volumes cover the dorsal-medial aspects of the TPCs, while the pterygoid air volumes extend forward to cover the ventral surface of the upper jaw-skull complex (Fig. 1A). This spatial arrangement suggests that the peribullar and pterygoid air volumes block sounds originating from the contralateral and dorsal-frontal directions, respectively, which we show by artificially manipulating the shape and orientation of model air volumes (Fig. S2B-C). In addition, other vestibular air volumes, such as the vestibular air sacs connected to the nasal passage, exist in live dolphins; these can vary in size and shape as the dolphins dive and recycle air to echolocate (Dormer, 1979). Although the premaxilla air sacs were not inflated in the CT scan images we used, simulation results from an artificially constructed premaxilla air volume suggest that its presence does not substantially alter the HRTFs (Fig. S2D). We propose that the pterygoid air volumes help stabilize the HRTFs by shielding against the otherwise potentially substantial effects of variable air sac morphology on the HRTFs.

HRTFs computed with the effects of shear waves in the bones, either from a full-head model (Fig. 8A) or from a reduced-head model consisting of only the non-fat soft tissues, TPCs, and the upper jaw-skull complex (Fig. 7B), show appreciable differences from HRTFs computed without shear wave effects, especially at lower frequencies (≲15 kHz). However, these differences largely disappear once the peribullar and pterygoid air volumes are included in the 3D model volume (Fig. 8C). The sound blocking effect of the air volumes likely explains the similarity between the HRTFs computed from the full-head model in this study and the HRTFs computed by modeling all bones as fluids in Aroyan, 2001 (see Fig. 10 and footnote 5 of that paper).

Our results suggest a limited role for the lower mandible in sound transmission, which corroborates with findings from a recent study (Wei et al., 2024) and differs from many prior studies. Specifically, the HRTFs computed with and without the lower mandible (Fig. 5E and Fig. 5B, respectively) showed only small differences in the form of altered interference patterns. This calls into question prior work arguing that the lower mandible serves as an effective pathway to conduct sound into the MFBs and then to the ears (Song et al., 2019, 2021) or that the lower mandible acts as a lens to focus the sound (Aroyan, 2001). Our observation is consistent with a recent modeling study that showed little change of displacement at the surface of periotic bones when the lower mandible was removed from the model volume (Wei et al., 2024). Nevertheless, the interference patterns shown in the HRTFs computed with the lower mandible included (Fig. 5E) do not rule out contributions to the HRTF interference patterns from flexural waves on the surface of the thin pan bone (Cranford et al., 2008). Indeed, solid plate structures can support flexural waves with a dispersive wave speed that varies with frequency (Graff, 1991). In addition, the periodic structure formed by teeth on the lower jaw has been proposed to function as the a directional beamformer (Dobbins, 2007), a directional bandstop filter (Dible et al., 2009), or an acoustic leaky wave antenna that selectively amplifies sound at different frequencies depending on the incident angle (Romero-Vivas and Leon-Lopez, 2025). Although the biological basis for these proposed mechanisms remains unclear, our results do not rule them out. Our shape representation focuses on gross anatomical structures and does not include fine details like individual teeth, and therefore may have limited our ability to capture mechanisms that depend on such structures. Further investigation is needed to fully describe the exact influence of the lower mandible and the teeth.

The HRTFs computed using the full-head model also are consistent with monaural hearing directivity derived from measured auditory evoked potential (AEP) in bottlenose dolphins. Most knowledge about hearing directivity in toothed whales comes from behavioral or physiological experiments in bottlenose dolphins and harbor porpoises, including AEPs measured with an electrode placed along the midline of the head (e.g., Accomando et al., 2020; Au and Moore, 1984; Kastelein et al., 2005; Taylor, 2013). In these experiments, however, the computed directivity derives from acoustic stimulation reaching both the left and right ears, so cannot be compared directly with our model predictions. Fortunately, monaural hearing directivity from bottlenose dolphins, measured by placing electrodes near the auditory meatus on the lateral head surface, reveals that the angle of maximum hearing sensitivity (the position with the lowest threshold) shifted gradually inward from 22.5° at 8-11.2 kHz to 0° at 90-128 kHz (Popov et al., 2006). While our models do not cover the full frequency range tested in these measurements, our computed HRTFs showed the same trend, where the region of maximum hearing sensitivity (highest HRTF values) moves toward the midline as frequency increases from 10 kHz to 50 kHz, with a slightly downward elevation angle (Fig. 4). Based on physical principles, we expect the HRTF predictions to be generally more reliable at lower frequencies, where sensitivity to fine shape details is less pronounced, than at higher frequencies. However, we caution that our modeling results also suggest that the predictions at lower frequencies are strongly influenced by the shapes of air volumes (Fig. 8; Sec. III. C), which are likely to vary significantly in live dolphins depending on behavioral context (e.g., diving depth).

The results presented in this study should be interpreted with the understanding that our predictions were influenced by the particular model parameterization choices we made, and that the model does not include the vibroacoustic process through which sound enters the middle ear. First, we assumed that all individual anatomical structures segmented out from the CT images (see Fig. 3) are homogeneous with the same material properties within the boundary of the structure. This approach differs from some previous studies that used regression relationships to infer sound speed and density of all soft tissues based on HU values from the CT scans (e.g., Song et al., 2019; Wei et al., 2024); however, we expect that small perturbations in the material properties would have modest effects on the general HRTF patterns. Second, the 3D representations of individual anatomical structures in the head were manually segmented from CT images (Sec. II. A), and the process involved some smoothing due to the resolution limits of the scans. Additionally, the CT images were from a living animal who could likely exert voluntary control over the exact shape of certain structures, such as the air volumes. Therefore, although the predicted HRTFs may not exactly match those of a dolphin, they are expected to capture the broad directional features and frequency-dependent trends. Third, we computed HRTFs at receiver locations right next to the tympanic bones, adjacent to the region where sound enters the middle ear (Hemilä et al., 1999, 2010). A bifurcated connection between the MFB and TPC through both the tympanic bone and the joint of the tympanic and periotic bones was described for multiple toothed whale species (Cranford et al., 2010), suggesting potentially complex propagation through the tympanic bones to the middle ear. To fully predict the sound reaching the cochlea and construct a complete model of the physics of dolphin hearing, one must account for this vibroacoustic coupling process and the response of the cochlea itself. Lastly, our model volume only included the dolphin head from the rostrum to the back of the skull. The influence of the rest of the dolphin body, in particular the air-filled lungs, may alter the exact HRTFs, especially when sound arrives from eccentric azimuthal angles (ϕ>90° and ϕ<-90° in the figures).

Intriguingly, our modeling results imply that evolution may have converged on the same solutions to tackle the challenge of creating strong directionality in both sound transmission and reception, which are reciprocal processes governed by the same physical principles. For sound reception, the MFBs act as acoustic lenses to guide sound waves toward the ears, counteracting the refraction caused by the higher sound speed of flesh (Fig. 5B-C). This mechanism functions in tandem with the peribullar and pterygoid air volumes, which isolate the ears from complex sound transmission through the bones and strongly shape the dolphin HRTFs (Fig. 5F and Fig. 8B-C). These mechanisms parallel those of the melon and vestibular air sacs in forming highly directional echolocation transmission beams (Cranford et al., 2014; Wei et al., 2016, 2017). However, we found the skull’s influence on the HRTFs to be more limited under the strong modulation of the air volumes, particularly at lower frequencies (⪅15 kHz) critical for dolphin communication (Fig. 8B-C), contrasting its important role in shaping the echolocation transmission beam (Aroyan et al., 1992; Cranford et al., 2014; Wei et al., 2016, 2017). These results suggest that the MFBs may have been adaptively selected to enhance localization of biosonar echoes, and the air volumes may have evolved to support sound localization in the frequency range of communication. However, as biological structures are typically shaped by multiple selective pressures, cross-species comparisons within a phylogenetic framework is needed to understand the potential evolutionary processes.

In this study, we focused on understanding the dominant sound transmission processes from the water to near the middle ear in the toothed whale head. We compared computationally predicted HRTFs to evaluate the relative importance of different anatomical structures to the overall propagation patterns, and utilized simplified analytical and numerical models to pinpoint the underlying physical processes. Our findings highlight convergent evolutionary solutions in toothed whale anatomy in creating strong directionality in the reciprocal sound transmission and reception processes, and our iterative modeling approach lays the groundwork for investigating the influence of anatomical variability across individuals and different toothed whale species. More broadly, development of dolphin HRTFs can inform the design of behavioral and neurophysiological study of dolphin auditory perception by providing an avenue to predict the acoustic scenes the animals are likely exposed to during experiments, to study how dolphins engage in spatial discrimination during different tasks. For example, listening for signs of prey or eavesdropping on other dolphin’s communication signals or echolocation returns may require using spatial cues to direct attentional resources via mechanisms similar to those found in terrestrial mammals. The HRTF models may also help us understand what is possible in terms of comparative cognition, and whether we can adapt dichotic listening paradigms, which rely on delivering sounds independently to right and left ears, in dolphin studies.

## Supplemental Materials

See supplementary material at [URL will be inserted by AIP] for the derivation of reflection and transmission coefficients through two- and three-layer media in Text S1 and Fig. S1, the influence of air volumes in the dolphin head in Fig. S2, and predicted HRTFs at both the left and right ears in Fig. S3-S6.

## Acknowledgments

We thank Arthur Cooper of Dolphins Plus in Florida, USA, for providing the bottlenose dolphin CT scan images. This work was supported by the Office of Naval Research (ONR) Multidisciplinary University Research Initiatives (MURI) Program grants N00014-18-1-2069, N00014-20-1-2709, and N00014-23-12065. The modeling used PSC Bridges-2 partitions at Pittsburgh Supercomputing Center through allocation BIO220174 from the Advanced Cyberinfrastructure Coordination Ecosystem: Services & Support (ACCESS) program, which is supported by National Science Foundation grants #2138259, #2138286, #2138307, #2137603, and #2138296. The funder had no role in study design, data collection and analysis, decision to publish, or preparation of the manuscript.

## Author Contributions

W.-J.L. conceived of the idea. Y.C. performed numerical modeling. W.-J.L. and Y.C. interpreted the results and wrote the manuscript. A.R. and J.K. provided the CT scan images. M.D.S. and B.S.-C. contributed to interpretation of results and editing the manuscript. All authors provided critical feedback and contributed to the final manuscript.

## Author Declarations

### Ethics Approval

The bottlenose dolphin was maintained by Dolphin Plus, Key Largo, Florida under the supervision of a full-time on-site veterinarian, who also oversaw the CT scan procedure.

### Data Availability

The data that support the findings of this study are available from the corresponding author upon reasonable request.

## References

Accomando, A. W., Mulsow, J., Branstetter, B. K., Schlundt, C. E., and Finneran, J. J. (2020). “Directional hearing sensitivity for 2–30 kHz sounds in the bottlenose dolphin (Tursiops truncatus),” J. Acoust. Soc. Am., 147, 388–398. doi:10.1121/10.0000557

Aroyan, J. L., Cranford, T. W., Kent, J., and Norris, K. S. (1992). “Computer modeling of acoustic beam formation in Delphinus delphis,” J. Acoust. Soc. Am., 92, 2539–2545. doi:10.1121/1.404424

Aroyan, J. L. (2001). “Three-dimensional modeling of hearing in Delphinus delphis,” J. Acoust. Soc. Am., 110, 3305–3318. doi:10.1121/1.1401757

Au, W. W. (1993). The sonar of dolphins, Springer Science & Business Media.

Au, W. W. L. (2015). “History of Dolphin Biosonar Research,” Acoustics today,.

Au, W. W. L., Kastelein, R. A., Benoit-Bird, K. J., Cranford, T. W., and McKenna, M. F. (2006). “Acoustic radiation from the head of echolocating harbor porpoises(Phocoena phocoena),” J. Exp. Biol., 209, 2726–2733. doi:10.1242/jeb.02306

Au, W. W. L., and Martin, S. W. (2012). “Why dolphin biosonar performs so well in spite of mediocre ‘equipment,’” IET Radar Sonar Amp Navig., 6, 566–575. doi:10.1049/iet-rsn.2011.0194

Au, W. W. L., and Moore, P. W. B. (1984). “Receiving beam patterns and directivity indices of the Atlantic bottlenose dolphin Tursiops truncatus,” J. Acoust. Soc. Am., 75, 255–262. doi:10.1121/1.390403

Au, W. W. L., and Pawloski, D. A. (1992). “Cylinder wall thickness difference discrimination by an echolocating Atlantic bottlenose dolphin,” J. Comp. Physiol. A Neuroethol. Sens. Neural. Behav. Physiol., 170, 41–47.

Au, W. W. L., and Snyder, K. J. (1980). “Long-range target detection in open waters by an echolocating Atlantic Bottlenose dolphin (Tursiops truncatus),” J. Acoust. Soc. Am., 68, 1077–1084. doi:10.1121/1.384993

Beaver, W. L., Dameron, D. H., and Macovski, A. (1977). “Ultrasonic Imaging with an Acoustic Lens,” IEEE Trans. Sonics Ultrason., 24, 235–243. Presented at the IEEE Transactions on Sonics and Ultrasonics. doi:10.1109/T-SU.1977.30937

Branstetter, B. K., Van Alstyne, K. R., Strahan, M. G., Tormey, M. N., Wu, T., Breitenstein, R. A., Houser, D. S., et al. (2020). “Spectral cues and temporal integration during cylinder echo discrimination by bottlenose dolphins (Tursiops truncatus),” J. Acoust. Soc. Am., 148, 614–626. doi:10.1121/10.0001626

Brill, R. L., Sevenich, M. L., Sullivan, T. J., Sustman, J. D., and Witt, R. E. (1988). “Behavioral Evidence for Hearing Through the Lower Jaw by an Echolocating Dolphin (tursiops Truncatus),” Mar. Mammal Sci., 4, 223–230. doi:10.1111/j.1748-7692.1988.tb00203.x

Coombs, E. J., Clavel, J., Park, T., Churchill, M., and Goswami, A. (2020). “Wonky whales: the evolution of cranial asymmetry in cetaceans,” BMC Biol., 18, 86. doi:10.1186/s12915-020-00805-4

Cranford, T. W., Amundin, M., and Norris, K. S. (1996). “Functional morphology and homology in the odontocete nasal complex: Implications for sound generation,” J. Morphol., 228, 223–285. doi:10.1002/(SICI)1097-4687(199606)228:3<223::AID-JMOR1>3.0.CO;2-3

Cranford, T. W., Krysl, P., and Amundin, M. (2010). “A New Acoustic Portal into the Odontocete Ear and Vibrational Analysis of the Tympanoperiotic Complex,” PLOS ONE, 5, e11927. doi:10.1371/journal.pone.0011927

Cranford, T. W., Krysl, P., and Hildebrand, J. A. (2008). “Acoustic pathways revealed: simulated sound transmission and reception in Cuvier’s beaked whale (Ziphius cavirostris),” Bioinspir. Biomim., 3, 016001. doi:10.1088/1748-3182/3/1/016001

Cranford, T. W., Trijoulet, V., Smith, C. R., and Krysl, P. (2014). “Validation of a vibroacoustic finite element model using bottlenose dolphin simulations: the dolphin biosonar beam is focused in stages,” Bioacoustics, 23, 161–194. doi:10.1080/09524622.2013.843061

Dible, S. A., Flint, J. A., and Lepper, P. A. (2009). “On the role of periodic structures in the lower jaw of the atlantic bottlenose dolphin (Tursiops truncatus),” Bioinspir. Biomim., 4, 015005. doi:10.1088/1748-3182/4/1/015005

Dobbins, P. (2007) “Dolphin sonar—modelling a new receiver concept,” Bioinspir. Biomim., 2, 19–29. doi: 10.1088/1748-3182/2/1/003

Dormer, K. J. (1979). “Mechanism of sound production and air recycling in delphinids: Cineradiographic evidence,” J. Acoust. Soc. Am., 65, 229–239. doi:10.1121/1.382240

Faran, J. J. (1951). “Sound Scattering by Solid Cylinders and Spheres,” J. Acoust. Soc. Am., 23, 405–418. doi:10.1121/1.1906780

Fedorov, A., Beichel, R., Kalpathy-Cramer, J., Finet, J., Fillion-Robin, J.-C., Pujol, S., Bauer, C., et al. (2012). “3D Slicer as an Image Computing Platform for the Quantitative Imaging Network,” Magn. Reson. Imaging, 30, 1323–1341. doi:10.1016/j.mri.2012.05.001

Graff, K. F. (1991). Wave Motion in Elastic Solids, Dover Publications, 688 pages. Retrieved from https://store.doverpublications.com/products/9780486667454

Gumerov, N. A., O’Donovan, A. E., Duraiswami, R., and Zotkin, D. N. (2010). “Computation of the head-related transfer function via the fast multipole accelerated boundary element method and its spherical harmonic representation,” J. Acoust. Soc. Am., 127, 370–386. doi:10.1121/1.3257598

Hemilä, S., Nummela, S., and Reuter, T. (1999). “A model of the odontocete middle ear,” Hear. Res., 133, 82–97. doi:10.1016/S0378-5955(99)00055-6

Hemilä, S., Nummela, S., and Reuter, T. (2010). “Anatomy and physics of the exceptional sensitivity of dolphin hearing (Odontoceti: Cetacea),” J. Comp. Physiol. A, 196, 165–179. doi:10.1007/s00359-010-0504-x

Houser, D. S., Finneran, J., Carder, D., Van Bonn, W., Smith, C., Hoh, C., Mattrey, R., et al. (2004). “Structural and functional imaging of bottlenose dolphin (Tursiops truncatus) cranial anatomy,” J. Exp. Biol., 207, 3657–3665. doi:10.1242/jeb.01207

Kastelein, R. A., Janssen, M., Verboom, W. C., and de Haan, D. (2005). “Receiving beam patterns in the horizontal plane of a harbor porpoise (Phocoena phocoena),” J. Acoust. Soc. Am., 118, 1172–1179. doi:10.1121/1.1945565

Ketten, D. R. (1994). “Functional analyses of whale ears: adaptations for underwater hearing,” Proc. Ocean., I/264-I/270 vol.1. Presented at the Proceedings of OCEANS’94. doi:10.1109/OCEANS.1994.363871

Ketten, D. R. (1997). “Structure and Function in Whale Ears,” Bioacoustics, 8, 103–135. doi:10.1080/09524622.1997.9753356

Knudsen, E. I., and Konishi, M. (1979). “Mechanisms of sound localization in the barn owl (*Tyto alba*),” J. Comp. Physiol., 133, 13–21. doi:10.1007/BF00663106

Krysl, P., and Cranford, T. W. (2016). “Directional Hearing and Head-Related Transfer Function in Odontocete Cetaceans,” In A. N. Popper and A. Hawkins (Eds.), Eff. Noise Aquat. Life II, Advances in Experimental Medicine and Biology, Springer, New York, NY, 583–587. doi:10.1007/978-1-4939-2981-8_70

Ladegaard, M., Mulsow, J., Houser, D. S., Jensen, F. H., Johnson, M., Madsen, P. T., and Finneran, J. J. (2019). “Dolphin echolocation behaviour during active long-range target approaches,” J. Exp. Biol., 222, jeb189217. doi:10.1242/jeb.189217

McCormick, J. G., Wever, E. G., Palin, J., and Ridgway, S. H. (1970). “Sound conduction in the dolphin ear,” J. Acoust. Soc. Am., 48, 1418–1428.

McCormick, J. G., Wever, E. G., Ridgway, S. H., and Palin, J. (1980). “Sound Reception in the Porpoise as it Relates to Echolocation,” In R.-G. Busnel and J. F. Fish (Eds.), Anim. Sonar Syst., Springer US, Boston, MA, pp. 449–467. doi:10.1007/978-1-4684-7254-7_18

McKenna, M. F., Cranford, T. W., Berta, A., and Pyenson, N. D. (2012). “Morphology of the odontocete melon and its implications for acoustic function,” Mar. Mammal Sci., 28, 690– 713. doi:10.1111/j.1748-7692.2011.00526.x

Mohl, B., Au, W. W. L., Pawloski, J., and Nachtigall, P. E. (1999). “Dolphin hearing: Relative sensitivity as a function of point of application of a contact sound source in the jaw and head region,” J. Acoust. Soc. Am., 105, 3421–3424. doi:10.1121/1.426959

Norris, K. S. (1968). “The evolution of acoustic mechanisms in odontocete cetaceans,” Evol. Environ.,.

Popov, V. V., Supin, A. Y., Klishin, V. O., and Bulgakova, T. N. (2006). “Monaural and binaural hearing directivity in the bottlenose dolphin: Evoked-potential study,” J. Acoust. Soc. Am., 119, 636–644. doi:10.1121/1.2141093

Romero-Vivas E., and Leon-Lopez, B. (2025) “Analogy of the dolphin jaw to a metamaterial leaky wave antenna for sound directional detection,” Journal of Sound and Vibration, 604, 118987. doi: 10.1016/j.jsv.2025.118987

Roston, R. A., and Roth, V. L. (2019). “Cetacean Skull Telescoping Brings Evolution of Cranial Sutures into Focus,” Anat. Rec., 302, 1055–1073. doi:10.1002/ar.24079

Song, Z., Zhang, J., Ou, W., Zhang, C., Dong, L., Dong, J., Li, S., et al. (2021). “Numerical-modeling-based investigation of sound transmission and reception in the short-finned pilot whale (Globicephala macrorhynchus),” J. Acoust. Soc. Am., 150, 225–232. doi:10.1121/10.0005518

Song, Z., Zhang, Y., Berggren, P., and Wei, C. (2017). “Reconstruction of the forehead acoustic properties in an Indo-Pacific humpback dolphin (Sousa chinensis), with investigation on the responses of soft tissue sound velocity to temperature,” J. Acoust. Soc. Am., 141, 681. doi:10.1121/1.4974861

Song, Z., Zhang, Y., Mooney, T. A., Wang, X., Smith, A. B., and Xu, X. (2019). “Investigation on acoustic reception pathways in finless porpoise (Neophocaena asiaorientalis sunameri) with insight into an alternative pathway,” Bioinspir. Biomim., 14, 016004. doi:10.1088/1748-3190/aaeb01

Taylor, K. A. (2013). Directional hearing and a head-related transfer function (HRTF) of a bottlenose dolphin (tursiops truncatus) University of Hawaii at Manoa. Retrieved from http://hdl.handle.net/10125/102027

Wei, C., Au, W. W. L., Song, Z., and Zhang, Y. (2016). “The role of various structures in the head on the formation of the biosonar beam of the baiji (Lipotes vexillifer),” J. Acoust. Soc. Am., 139, 875–880. doi:10.1121/1.4941780

Wei, C., Au, W. W. L., Ketten, D. R., Song, Z., and Zhang, Y. (2017). “Biosonar signal propagation in the harbor porpoise’s (Phocoena phocoena) head: The role of various structures in the formation of the vertical beam,” J. Acoust. Soc. Am., 141, 4179–4187. doi:10.1121/1.4983663

Wei, C., Erbe, C., Smith, A. B., and Yang, W.-C. (2024). “Validated 3D finite-element model of the Risso’s dolphin (Grampus griseus) head anatomy demonstrates gular sound reception and channelling through the mandibular fats,” Bioinspir. Biomim., 19, 056025. doi:10.1088/1748-3190/ad7344

