## Supplemental text and figures for "Head-related transfer function predictions reveal dominant sound transmission mechanisms in a dolphin head"

### Supplemental Materials

#### 1 Reflection and transmission through two- and three-layer media

This section provides analytical solutions for the reflection ($R$) and transmission ($T$) coefficients of waves propagating across two or three layers of infinite and finite media (Fig. S1) used for interpreting the HRTF results. Both pressure and shear waves are considered in the general solution derived in Aki and Richards (2002).

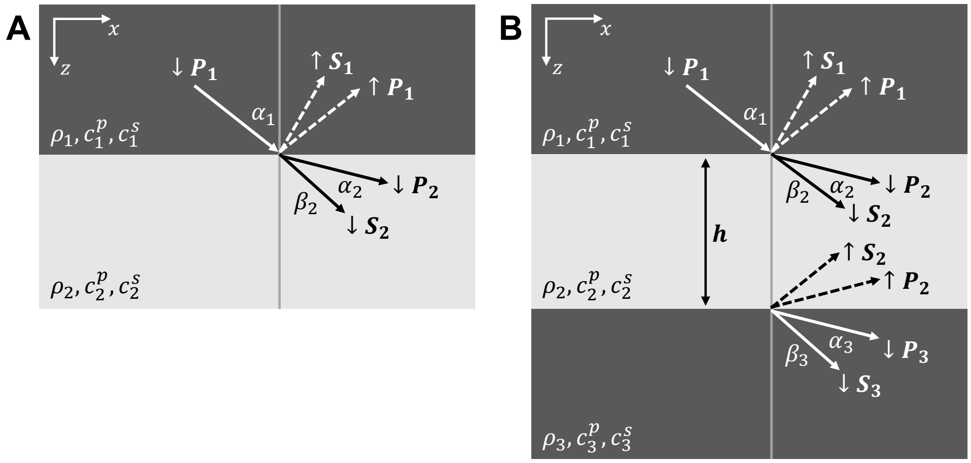


Figure S1: Schematics of the two-layer (A) and three-layer (B) setup with both pressure and shear waves considered in all media. Here, $\rho_n$, $c_n^p$, and $c_n^s$ denote the density, pressure wave speed, and shear wave speed in medium $n$, respectively. $P_n$ and $S_n$ denote the pressure wave and shear wave in medium $n$, respectively.

In the two-layer case (Fig. S1A), the plane $z = 0$ is stress-free and the reflection and transmission coefficients can be solved using the four boundary conditions of continuity of displacement and traction at the boundary between media 1 and 2. Specifically, by expressing the pressure and shear waves using the potentials $\phi$ and $\psi$, respectively, and using symbols $\downarrow$ and $\uparrow$ to describe downward and upward propagating waves, respectively, the reflected and transmitted pressure and shear waves are given by:

$$
\begin{aligned}
\downarrow \phi_1 &= \frac{c_1^p}{i\omega} \exp\left(i\omega(px + \frac{\cos\alpha_1}{c_1^p} z - t)\right) \\
\uparrow \phi_1 &= A_1 \frac{c_1^p}{i\omega} \exp\left(i\omega(px - \frac{\cos\alpha_1}{c_1^p} z - t)\right) \\
\uparrow \psi_1 &= A_2 \frac{c_1^s}{i\omega} \exp\left(i\omega(px - \frac{\cos\beta_1}{c_1^s} z - t)\right) \\
\downarrow \phi_2 &= A_3 \frac{c_2^p}{i\omega} \exp\left(i\omega(px + \frac{\cos\alpha_2}{c_2^p} z - t)\right) \\
\downarrow \psi_2 &= -A_4 \frac{c_2^s}{i\omega} \exp\left(i\omega(px + \frac{\cos\beta_2}{c_2^s} z - t)\right)
\end{aligned} \tag{1}
$$

where

$$
p = \frac{\sin\alpha_1}{c_1^p} = \frac{\sin\beta_1}{c_1^s} = \frac{\sin\alpha_2}{c_2^p} = \frac{\sin\beta_2}{c_2^s} \tag{2}
$$

through Snell's Law, and $A_1$, $A_2$, $A_3$, and $A_4$ are four unknown coefficients.

The displacement vectors and traction vectors for the pressure and shear waves can be expressed as:

$$\begin{aligned}
u_\phi &= (i\omega p\phi, 0, \frac{\partial \phi}{\partial z}),\\
u_\psi &= \left(-\frac{\partial \psi}{\partial z}, 0, i\omega p\psi\right),\\
\tau_\phi &= \left(2i\omega p\rho(c^s)^2\frac{\partial \phi}{\partial z}, 0, -\rho\left(1-2(c^s p)^2\right)\omega^2\phi\right), \text{ and}\\
\tau_\psi &= \left(\rho\left(1-2(c_s p)^2\right)\omega^2\psi, 0, 2i\omega p\rho(c^s)^2\frac{\partial \psi}{\partial z}\right),
\end{aligned} \tag{3}$$

respectively. Substituting the potentials to satisfy the boundary conditions for the continuity of $u_x$, $u_z$, $\tau_{zx}$, and $\tau_{zz}$ yields the following solutions for the unknown coefficients

$$\begin{bmatrix} A_1 \\ A_2 \\ A_3 \\ A_4 \end{bmatrix} = \begin{bmatrix} \sin\alpha_1 & \cos\beta_1 & -\sin\alpha_2 & -\cos\beta_2 \\ -\cos\alpha_1 & \sin\beta_1 & -\cos\alpha_2 & \sin\beta_2 \\ -2\rho_1 {c_1^s}^2 p\cos\alpha_1 & -\rho_1(1-2{c_1^s}^2p^2)c_1^s & -2\rho_2 {c_2^s}^2 p\cos\alpha_2 & -\rho_2(1-2{c_2^s}^2p^2)c_2^s \\ \rho_1(1-2{c_1^s}^2p^2)c_1^p & -2\rho_1 {c_1^s}^2 p\cos\beta_1 & -\rho_2(1-2{c_2^s}^2p^2)c_2^p & 2\rho_2 {c_2^s}^2 p\cos\beta_2 \end{bmatrix}^{-1} \times \begin{bmatrix} -\sin\alpha_1 \\ -\cos\alpha_1 \\ -2\rho_1 {c_1^s}^2 p\cos\alpha_1 \\ -\rho_1(1-2{c_1^s}^2p^2)c_1^p \end{bmatrix} \tag{4}$$

In the special scenario in which media 1 and 2 are both fluid media that support only pressure waves, the above reduces to

$$\begin{bmatrix} A_1 \\ A_3 \end{bmatrix} = \begin{bmatrix} -\cos\alpha_1 & -\cos\alpha_2 \\ \rho_1 c_1^p & -\rho_2 c_2^p \end{bmatrix}^{-1} \times \begin{bmatrix} -\cos\alpha_1 \\ -\rho_1 c_1^p \end{bmatrix}, \tag{5}$$

for which the reflection and transmission coefficients at the boundary are

$$\begin{aligned}
R = A_1 &= \frac{m\cos\alpha_1 - n\sqrt{1-\sin^2\alpha_1/n^2}}{m\cos\alpha_1 + n\sqrt{1-\sin^2\alpha_1/n^2}}\\
T = \frac{\rho_2}{\rho_1}\frac{c_2^p}{c_1^p}A_3 &= \frac{2m\cos\alpha_1}{m\cos\alpha_1 + n\sqrt{1-\sin^2\alpha_1/n^2}}
\end{aligned} \tag{6}$$

where $n = c_1^p/c_2^p$ and $m = \rho_2/\rho_1$. For the interface between water ($c_1^p = 1500$ m/s, $\rho_1 = 1000$ kg/m$^3$) and air ($c_2^p = 343$ m/s, $\rho_2 = 1.2$ kg/m$^3$), $R \approx 1$ and $T \approx 0$.

In the more complex case where a finite layer is sandwiched between two infinite media (Fig. S1B), by following the same procedure as the above using the eight boundary conditions for the continuity of displacement and traction at the boundaries $z = 0$ and $z = h$, the unknown coefficients that fully characterize all waves in the three media are

$$
\begin{bmatrix} A_1 \\ A_2 \\ A_3 \\ A_4 \\ A_5 \\ A_6 \\ A_7 \\ A_8 \end{bmatrix}
=
\begin{bmatrix}
\sin\alpha_1 & \cos\beta_1 & -\sin\alpha_2 & -\cos\beta_2 & -\sin\alpha_2 & -\cos\beta_2 & 0 & 0 \\
-\cos\alpha_1 & \sin\beta_1 & -\cos\alpha_2 & \sin\beta_2 & \cos\alpha_2 & -\sin\beta_2 & 0 & 0 \\
-2\rho_1 {c_1^s}^2 p\cos\alpha_1 & -\rho_1(1-2{c_1^s}^2p^2)c_1^s & -2\rho_2 {c_2^s}^2 p\cos\alpha_2 & -\rho_2(1-2{c_2^s}^2p^2)c_2^s & 2\rho_2 {c_2^s}^2 p\cos\alpha_2 & \rho_2(1-2{c_2^s}^2p^2)c_2^s & 0 & 0 \\
\rho_1(1-2{c_1^s}^2p^2)c_1^p & -2\rho_1 {c_1^s}^2 p\cos\beta_1 & -\rho_2(1-2{c_2^s}^2p^2)c_2^p & 2\rho_2 {c_2^s}^2 p\cos\beta_2 & -\rho_2(1-2{c_2^s}^2p^2)c_2^p & 2\rho_2 {c_2^s}^2 p\cos\beta_2 & 0 & 0 \\
0 & 0 & \sin\alpha_2 e^{i\omega h\frac{\cos\alpha_2}{c_2^p}} & \cos\beta_2 e^{i\omega h\frac{\cos\beta_2}{c_2^s}} & \sin\alpha_2 e^{-i\omega h\frac{\cos\alpha_2}{c_2^p}} & \cos\beta_2 e^{-i\omega h\frac{\cos\beta_2}{c_2^s}} & -\sin\alpha_3 e^{i\omega h\frac{\cos\alpha_3}{c_3^p}} & -\cos\beta_3 e^{i\omega h\frac{\cos\beta_3}{c_3^s}} \\
0 & 0 & \cos\alpha_2 e^{i\omega h\frac{\cos\alpha_2}{c_2^p}} & -\sin\beta_2 e^{i\omega h\frac{\cos\beta_2}{c_2^s}} & -\cos\alpha_2 e^{-i\omega h\frac{\cos\alpha_2}{c_2^p}} & \sin\beta_2 e^{-i\omega h\frac{\cos\beta_2}{c_2^s}} & -\cos\alpha_3 e^{i\omega h\frac{\cos\alpha_3}{c_3^p}} & \sin\beta_3 e^{i\omega h\frac{\cos\beta_3}{c_3^s}} \\
0 & 0 & 2\rho_2 {c_2^s}^2 p\cos\alpha_2 e^{i\omega h\frac{\cos\alpha_2}{c_2^p}} & \rho_2(1-2{c_2^s}^2p^2)c_2^s e^{i\omega h\frac{\cos\beta_2}{c_2^s}} & -2\rho_2 {c_2^s}^2 p\cos\alpha_2 e^{-i\omega h\frac{\cos\alpha_2}{c_2^p}} & -\rho_2(1-2{c_2^s}^2p^2)c_2^s e^{-i\omega h\frac{\cos\beta_2}{c_2^s}} & -2\rho_3 {c_3^s}^2 p\cos\alpha_3 e^{i\omega h\frac{\cos\alpha_3}{c_3^p}} & -\rho_3(1-2{c_3^s}^2p^2)c_3^s e^{i\omega h\frac{\cos\beta_3}{c_3^s}} \\
0 & 0 & \rho_2(1-2{c_2^s}^2p^2)c_2^p e^{i\omega h\frac{\cos\alpha_2}{c_2^p}} & -2\rho_2 {c_2^s}^2 p\cos\beta_2 e^{i\omega h\frac{\cos\beta_2}{c_2^s}} & \rho_2(1-2{c_2^s}^2p^2)c_2^p e^{-i\omega h\frac{\cos\alpha_2}{c_2^p}} & -2\rho_2 {c_2^s}^2 p\cos\beta_2 e^{-i\omega h\frac{\cos\beta_2}{c_2^s}} & -\rho_3(1-2{c_3^s}^2p^2)c_3^p e^{i\omega h\frac{\cos\alpha_3}{c_3^p}} & 2\rho_3 {c_3^s}^2 p\cos\beta_3 e^{i\omega h\frac{\cos\beta_3}{c_3^s}}
\end{bmatrix}^{-1}
\times
\begin{bmatrix} -\sin\alpha_1 \\ -\cos\alpha_1 \\ -2\rho_1 {c_1^s}^2 p\cos\alpha_1 \\ -\rho_1(1-2{c_1^s}^2p^2)c_1^p \\ 0 \\ 0 \\ 0 \\ 0 \end{bmatrix}
\tag{7}
$$

The transmission and reflection coefficients in Fig. 7C-D were computed by numerically inverting the matrix above and using

$$
\begin{aligned}
R &= A_1 \\
T &= \frac{\rho_3(1-2{c_3^s}^2p^2)c_3^p e^{i\omega h\frac{\cos\alpha_3}{c_3^p}}}{\rho_1(1-2{c_1^s}^2p^2)c_1^p} A_7,
\end{aligned}
\tag{8}
$$

where $T$ characterizes the transmission from medium 1 to medium 3. Note that $R$ and $T$ are both functions of $\omega h$, and therefore functions of $k_n h$, where $k_n$ is the wavenumber of medium $n$.

In the simplified scenario in which all three media only support pressure waves, the reflection coefficient has the closed-form expression (Jensen et al., 2011)

$$
R = \frac{Z_2(Z_3 - Z_1) - i(Z_2^2 - Z_1 Z_3)\tan\delta_2}{Z_2(Z_3 + Z_1) - i(Z_2^2 + Z_1 Z_3)\tan\delta_2}
\tag{9}
$$

where $Z_n = \frac{\rho_n c_n^p}{cos i_n}$, $k = 1, 2, 3$ and $\delta_2 = k_2 h\cos\alpha_2$.

By setting $Z_1 = Z_3$ (Fig. 7C) and noting that $\tan\delta_2 = \tan(k_2 h\cos\alpha_2) = \tan N\pi = 0$, where $N$ is an integer, $R = 0$ when $k_2 h\cos\alpha_2 = N\pi$. Using Snell's Law $\sin\alpha_1/c_1^p = \sin\alpha_2/c_2^p$ and relating the wavenumbers $k_2 = k_1 c_1^p/c_2^p$, we show that

$$
R = 0 \text{ when } k_1 h = N\pi\frac{c_2^p}{c_1^p}\frac{1}{\sqrt{1-(\frac{c_2^p}{c_1^p}\sin\alpha_1)^2}}.
\tag{10}
$$

These correspond to the $T = 1$ branches in Fig. 7C. For example, with pressure wave speeds $c_1^p = 1650$ m/s and $c_2^p = 3380$ m/s, incident angle $\alpha_1 = 0$, and $N = 1$, $T = 1$ for $k_1 h = 6.43$.

#### 2 Influence of air volumes

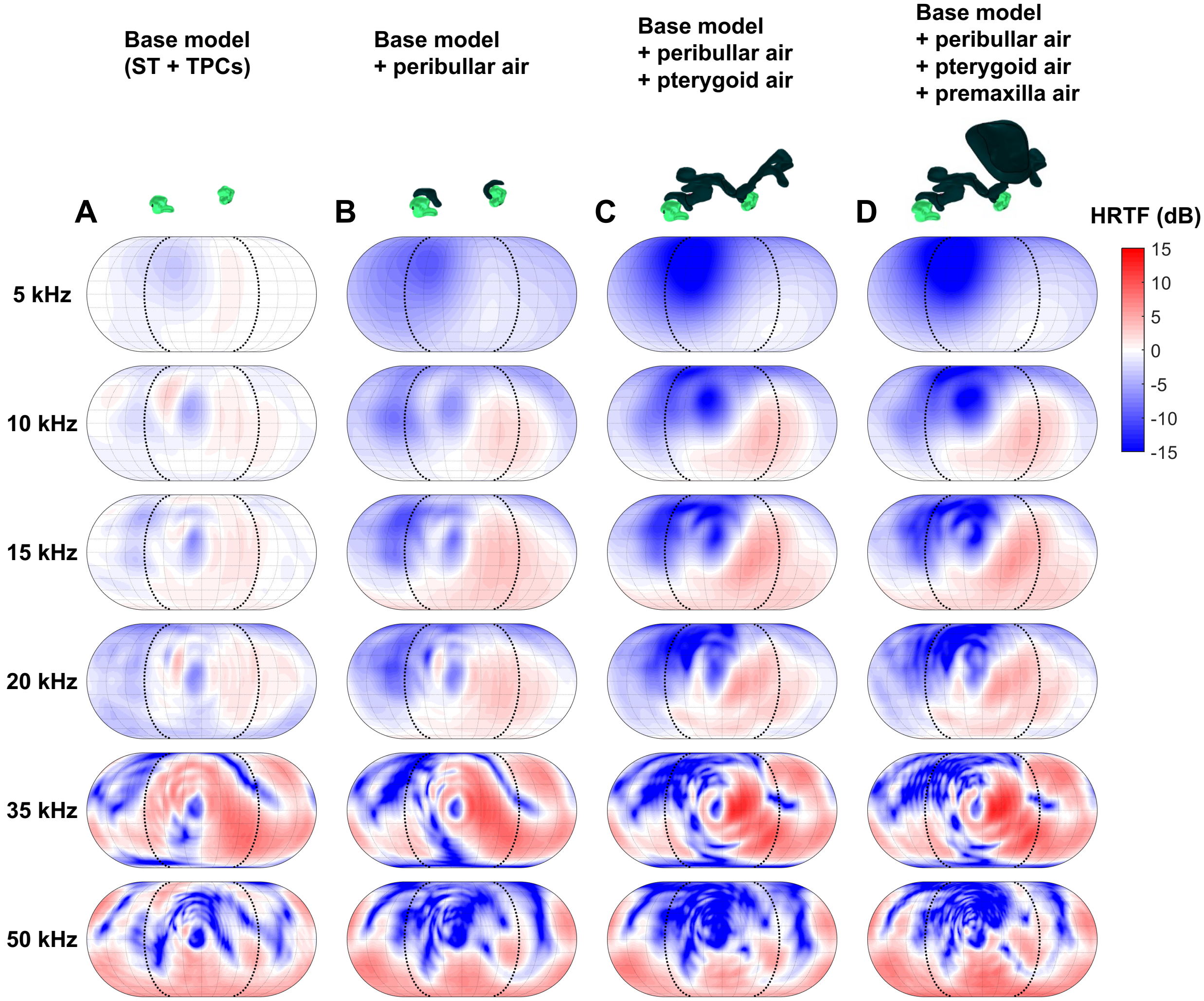


Figure S2: HRTFs computed using (A) the base model (non-fat soft tissues and the TPCs) and a series of reduced head model consisting of the non-fat soft tissues, TPCs, and different combinations of air volumes: (B) peribullar air volumes; (C) peribullar and pterygoid air volumes; and (D) peribullar, pterygoid, and premaxilla air volumes. Comparison across panels A-C show the peribullar air volumes block sounds from the contralateral direction and the additional effects of the pterygoid air volumes. Comparison between panels C-D show that the minimal influence of the premaxilla air volume with the peribullar and pterygoid air volumes forming an effective barrier to the ear.

#### 3 HRTFs at the left and right ears

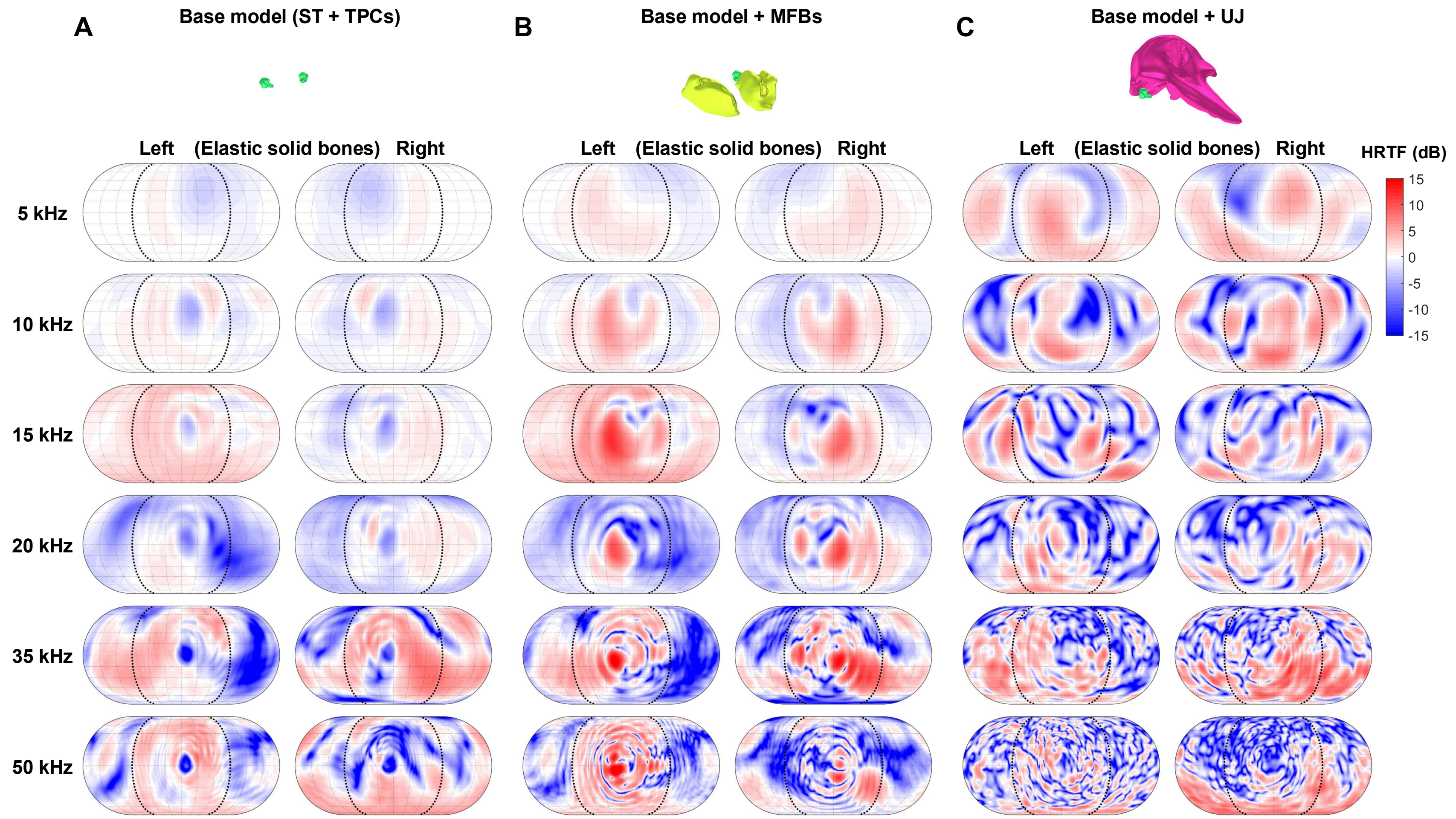


Figure S3: HRTFs at the left and right ears computed using (A) the base model (non-fat soft tissues and the TPCs), (B) the base model and the MFBs, and (C) the base model and the upper jaw-skull complex. The right columns of panels A, B, and C are identical to Fig. 5B, Fig. 5C, and Fig. 5D. All bones are modeled as elastic solids.

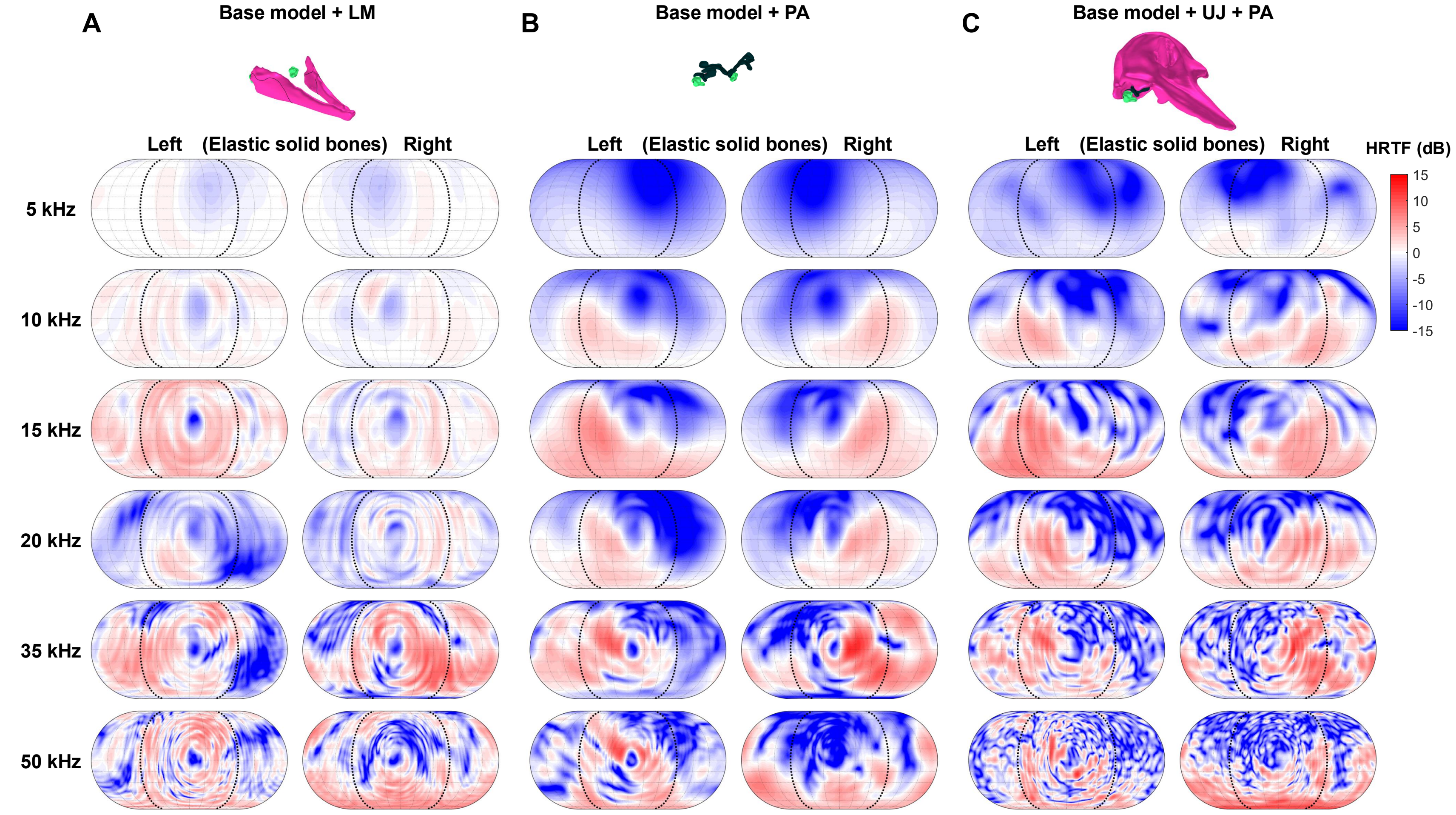


Figure S4: HRTFs at the left and right ears computed using (A) the base model and the lower mandible, (B) the base model and the peribullar and pterygoid air volumes, and (C) the base model, the upper jaw-skull complex, and the peribullar and pterygoid air volumes. The right columns of panels A, B, and C are identical to Fig. 5E, Fig. 5F, and the left column of Fig. 8C. All bones are modeled as elastic solids.

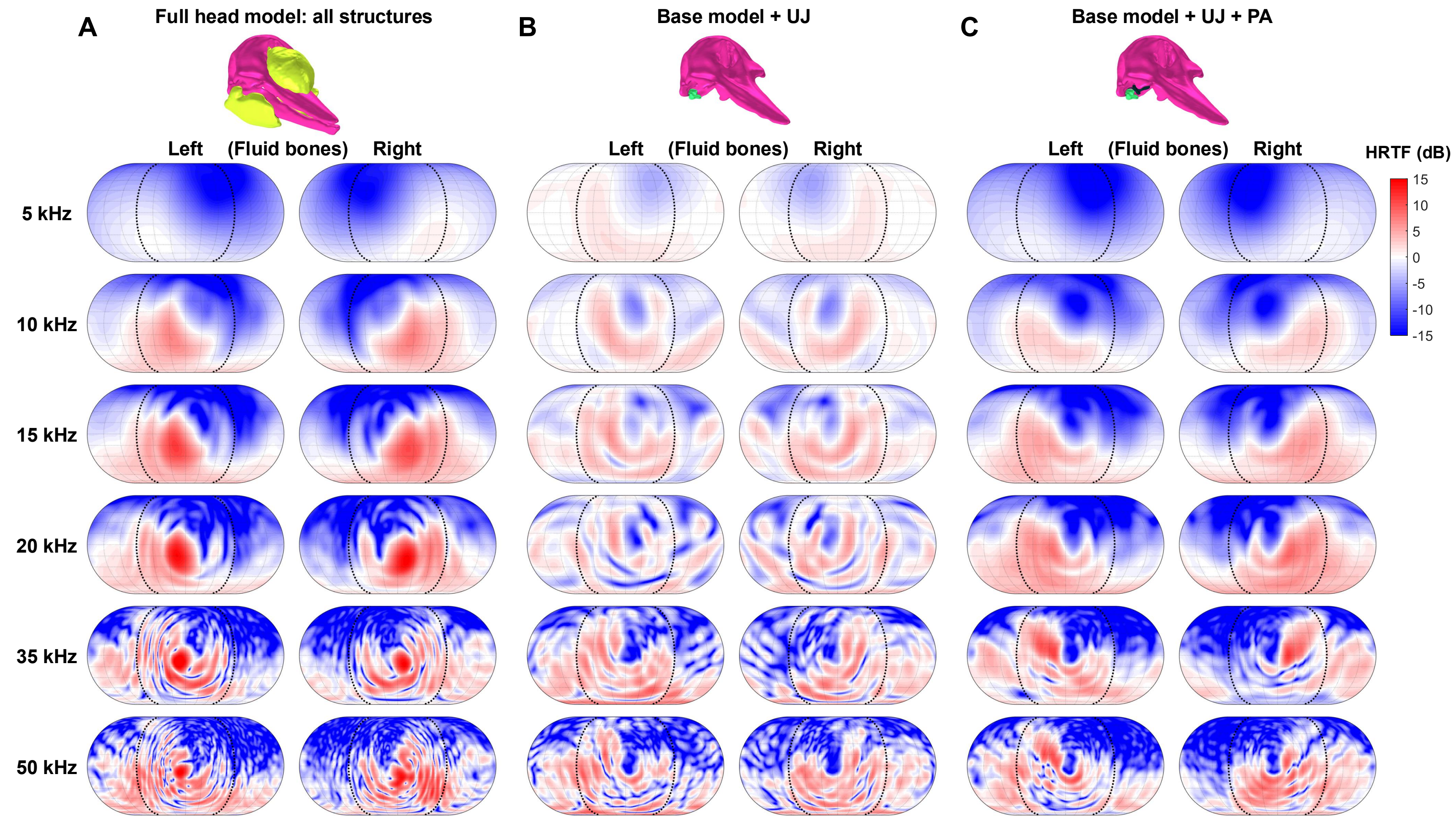


Figure S5: HRTFs at the left and right ears computed using (A) the full-head model, (B) the base model and the upper jaw-skull complex, and (C) the base model, the upper jaw-skull complex, and the peribullar and pterygoid air volumes. The right columns of panels A, B, and C are identical to the right columns of Fig. 8A, Fig. 8B, and Fig. 8C. All bones are modeled as fluids.

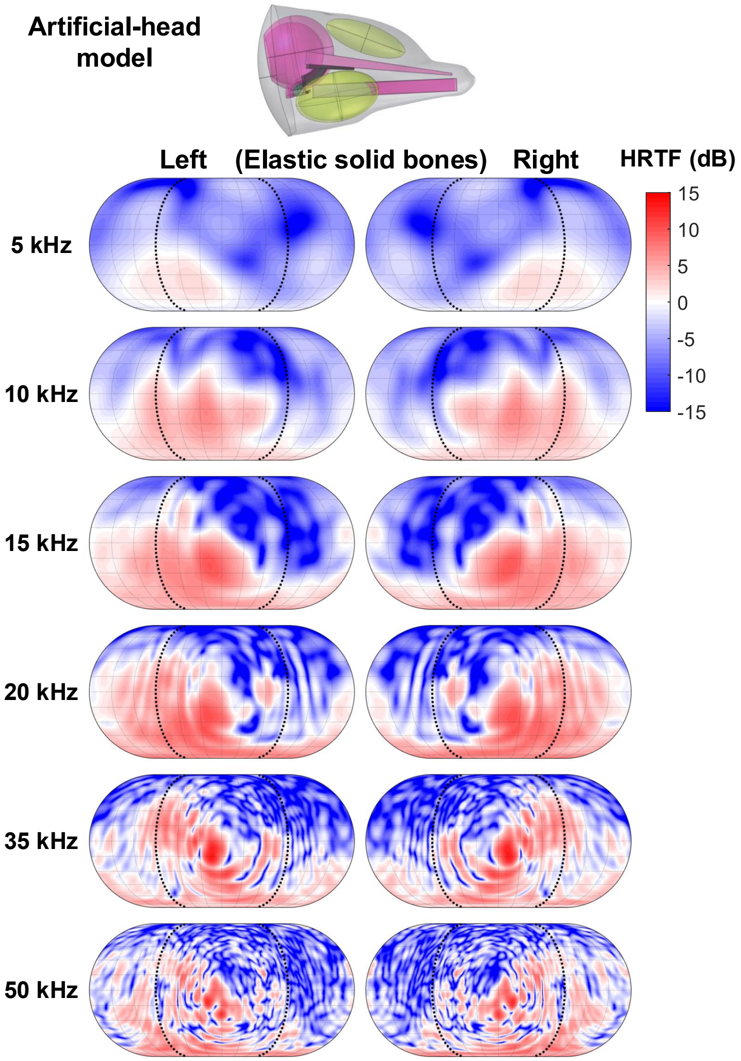


Figure S6: HRTFs at the left and right ears computed using the artificial-head model. The right column is identical to Fig. 9B. All bones are modeled as elastic solids.

### References


Aki, K., and Richards, P. G. (2002). "Quantitative Seismology, 2nd Ed.," University Science Books, Sausalito. p.132 equations 5.21-22.

Jensen, F. B., Kuperman, W. A., Porter, M. B., and Schmidt, H. (2011). Computational Ocean Acoustics, Modern Acoustics and Signal Processing, Springer, New York, NY. p.47 equation 1.69.
